# Recurrent drought increases grassland community seasonal synchrony

**DOI:** 10.1101/2024.01.29.577778

**Authors:** Lena M. Müller, Michael Bahn, Maximillian Weidle, Georg Leitinger, Dina in ‘t Zandt

## Abstract

1. Climate change increases the frequency and severity of drought events with strong repercussions on grassland ecosystems. While the effects of single drought events on ecosystem structure and functioning are well understood, it is largely unknown whether and how drought frequency modifies ecosystem responses to drought.

2. Here, we assessed how the increase in frequency of severe, annual summer drought impacted grassland communities. We examined these effects in a species-rich sub-alpine mountain meadow with a drought frequency of one, three, and 13 years, as well as ambient conditions.

3. We found that high drought frequency increased seasonal plant community synchrony through a reduction in species richness, a shift of plant functional groups, a loss of early-seasonal plant species, and the constrained establishment of seedlings throughout the growing season. These changes were associated with a decreased fraction of biomass as drought frequency increased.

4. Furthermore, we show that negative drought effects were enhanced with an increasing drought frequency, and that negative drought effects on plant communities outweighed the weak adaptive effects of species.

5. Synthesis. We conclude that single and low-frequency drought studies may not adequately predict longer-term changes in our rapidly shifting climate. With the ongoing increase in drought frequency due to climate change, we predict that grassland plant communities will increase in seasonal synchrony. We suggest that this increase in synchrony will leave ecosystems highly vulnerable to future disturbances, because asynchrony is a critical component of stability. Moreover, given the weak adaptive effects of plant species to long-term recurrent drought, we conclude that plant communities are unlikely to be able to adapt to the rapid increase in recurrent drought events.

## Introduction

Grasslands cover 40 % of the Earth’s surface and are crucial for carbon and nutrient cycling, biodiversity conservation, and ecosystem services (Petermann & Buzhdygan 2021; Blair *et al*. 2014; Dixon *et al*. 2014). However, up to 49 % of the world’s grasslands are currently degraded, with the climate crisis predicted to exacerbate this number (Bardgett *et al*. 2021; Gibbs & Salmon 2015). Among the detrimental climate scenarios, the increase in frequency and severity of drought events stand out as particularly damaging (IPCC 2023, 2021; Wang *et al*. 2021; Spinoni *et al*. 2018). Drought events alter grassland nutrient cycling, disrupt interactions between species, result in species extinction, and therewith cause severe disruptions in ecosystem structure and function (Smith *et al*. 2024; Oram *et al*. 2023; Bardgett & Caruso 2020; Ingrisch *et al*. 2020; Tello-García *et al*. 2020). In the long-term, the predicted increase in the frequency of drought events therefore threatens to irrevocably change grassland ecosystems, including the fundamental services and biodiversity they conserve (Müller & Bahn 2022; Xu *et al*. 2021; Schirpke *et al*. 2019). At the same time, potential adaptations of species and communities may dampen these detrimental effects to some extent (Müller & Bahn 2022; Canarini *et al*. 2021; Coleman & Wernberg 2020). This occurs for example through drought-induced legacy effects that alter ecosystem structure and function after drought has subsided, and which results in plant communities being able to cope better with a subsequent drought event (Müller & Bahn 2022). Yet, while the effects of single drought events are comparatively well studied, our understanding of the effects of a long-term increase in drought frequency on grassland community structure and function remains limited.

Multiple interlinked drought-induced changes, for example in plant species composition, diversity, complementarity, as well as species loss may have detrimental effects by affecting the **synchrony** of grassland communities (He *et al*. 2022; Muraina *et al*. 2021; De Mazancourt *et al*. 2013). Community synchrony refers to the degree to which populations of different species within a community fluctuate in tandem. High community synchrony indicates that populations of different species increase and decrease at the same points in time, while low synchrony (or asynchrony) indicates that species populations fluctuate independently of each other (Loreau & De Mazancourt 2013). With greater synchrony between years, species communities exhibit increased similarity in their response to environmental changes, the swiftness of their responses, or their functional complementarity throughout various seasons (Loreau & De Mazancourt 2013). Single drought was found to increase the synchrony of plant communities between years (Zhang *et al*. 2019), similarly as recurrent drought of three (He *et al*. 2022) and four years (Muraina *et al*. 2021). These changes in grassland community synchrony likely have detrimental effects on long-term ecological functioning, especially under a future climate with a high drought frequency. Increases in grassland community synchrony between years has been shown to decrease community stability (Wilcox *et al*. 2017; Xu *et al*. 2015; Loreau & De Mazancourt 2013). This is because asynchronous communities stabilise ecosystems through different responses to disturbances (De Mazancourt *et al*. 2013). For drought disturbances, species asynchrony was suggested to be the dominant driver to stabilise community productivity under drought (Luo *et al*. 2023b; Muraina *et al*. 2021; Craven *et al*. 2018). Community synchrony between years is thus influenced by drought and is suggested to shape a communities’ response to subsequent droughts by decreasing its stability. However, besides changing community synchrony *between* seasons, drought is also likely to affect community synchrony *within* the growing season. Therewith, it might be a critical mechanism in shaping responses to recurrent drought. However, we currently lack evidence on the effects of recurrent drought on community *seasonal* synchrony. In the presence of more frequent droughts, we hypothesize that the plant community will become more synchronous within the season.

Drought events alter plant community composition, phenology, and productivity (Smith *et al*. 2024; Müller & Bahn 2022). Drought leads to a decrease in plant community **productivity**, by limiting plant growth through reduced water availability, or by mortality of species under severe drought conditions (Sippel *et al*. 2018; Frank *et al*. 2015). Additionally, after a drought event, soil rewetting results in a temporarily increased availability of nitrogen (Ingrisch *et al*. 2018; Karlowsky *et al*. 2018). This surge in nitrogen supports the recovery and, in some cases, leads to a short-term overshoot in productivity (Griffin-Nolan *et al*. 2018; Hofer *et al*. 2016), which can compensate for the lower plant community productivity during the drought event (Hahn *et al*. 2021). This response is mainly based on fast-growing grass species, which makes the community sensitive to changes in composition post drought (Mackie *et al*. 2019; Stampfli *et al*. 2018). An important issue is that drought-induced changes in productivity and composition in grasslands may take a long time to come back to pre-disturbance level (Müller & Bahn 2022). If a subsequent drought falls within this recovery timespan of a previous drought, it may lead to constantly impaired ecosystems (Schwalm *et al*. 2017), and consequently to a more profound effect on productivity under a high frequency of drought (Xu *et al*. 2021). Hence, a potential accumulation of effects can emerge when multiple droughts occur annually. However, as recurrent drought events are typically studied in two-year experiments and only rarely extend beyond this timeframe, we lack critical knowledge on the productivity loss that builds up under a high drought frequency (Knapp *et al*. 2023; Hoover *et al*. 2018).

Drought shifts plant community **composition** towards more slower growing and more drought adapted-species (Wilcox *et al*. 2021). For example, drought may lead to a dominance of grasses over forbs (Xu *et al*. 2021; De Boeck *et al*. 2018; Hoover *et al*. 2014). This may be driven by a higher post-drought resource use of grasses compared with the conservative resource use of forbs (Stampfli *et al*. 2018) or by a more efficient resource acquisition of water of grasses compared with forbs (Tello-García *et al*. 2020). Furthermore, drought-induced changes in plant community composition are driven by a reduction in plant species performance and an increase in mortality of plants both during and after a drought event (Sippel *et al*. 2018; Frank *et al*. 2015). In the long term, plant compositional shifts due to high drought frequency are likely to relate additionally to drought-induced changes in plant reproductive effects, e.g. reduction in reproductive shoots, seeds (Zeiter *et al*. 2016), belowground bud density (Qian *et al*. 2022), as well as altered seed banks (Basto *et al*. 2018), and regeneration from seed (Stampfli & Zeiter 2004). Through these drought-induced changes in plant species composition, recurrent drought events may in the long-term result in permanent loss of plant species with potentially irreversible consequences for the functioning of plant communities. Furthermore, under recurrent drought, plant species also have the capacity to adapt (Müller & Bahn 2022), which dampens drought effects under a higher drought frequency (Backhaus *et al*. 2014; Walter *et al*. 2011). Therefore, a high drought frequency is expected to result in plant communities consisting of drought-tolerant species, a dominance of grasses, plant species profiting from the recovery dynamics of drought events, as well as plant species that adapt to recurrent drought conditions. However, due to the lack of studies investigating high drought frequency, long-term compositional shifts might have been overlooked to date.

Drought alters **phenology** by shifting the beginning or end of the growing season (Yuan *et al*. 2023; Ge *et al*. 2022; Rihan *et al*. 2022; Wang *et al*. 2022; Ji *et al*. 2021; Zeng *et al*. 2021; Yuan *et al*. 2020; Berwaers *et al*. 2019; Kang *et al*. 2018) and advancing end-of-season leaf senescence (Hoover *et al*. 2021; Berwaers *et al*. 2019; Peng *et al*. 2019; Kang *et al*. 2018). Furthermore, drought alters species flowering phenology by advancing the flowering date and extending the flowering length, whereby effects differ strongly between species (Castillioni *et al*. 2022; Vorkauf *et al*. 2021; Forrest & Miller-Rushing 2010; Jentsch *et al*. 2009). Hence, recurrent drought is expected to move the duration and timing of the growing season and flowering with effects on community composition, though the directions and magnitudes of these changes under recurrent drought are still poorly understood.

We lack a profound understanding of the implications of a high frequency of drought events on the temporal shifts in community composition, productivity, and phenology. Here, we assessed how the increase in frequency of severe summer drought affected grassland community structure, function, and seasonal dynamics. We examined this in a species-rich sub-alpine mountain meadow with increasing annual summer drought frequencies of up to 13 years. We quantified species abundances in the field throughout the growing season and sampled plant species-specific biomass and necromass at peak drought and drought recovery. We asked the following questions: How did an increase in drought frequency affect (1) plant community productivity, species richness, and flower production during peak drought and recovery, respectively, as well as (2) plant community seasonal synchrony. We examined whether these patterns were underlain by shifts in (3) seedling establishment (4), plant community composition, (5) plant functional groups, and (6) plant community traits. Taken together, we present the effects of a high frequency of drought events on plant communities and discuss underlying mechanisms.

## Material and Methods

### Study Site

The studied sub-alpine grassland is located in the Austrian Central Alps in the Stubai valley at 1820 m a.s.l. (47°70 45″N, 11°180 20″E). The site is exposed southeast with an inclination of ∼ 20° and has an average annual temperature of 3 °C and an annual precipitation of 1100 mm. The soil type is a dystric cambisol (top soil pH = 5.5). The vegetation community is classified as *Trisetetum flavescentis*, species-rich with on average 49 species per m^2^ and consists largely of perennial grasses and forbs. The growing season lasts from the end of April until mid-October. The sub-alpine grassland is manured every 2-3 years, mown once in early August at peak biomass, and sporadically grazed in spring and autumn. The studied area, however, has not been fertilised and is excluded from grazing since 2007. For further details on the site see Ingrisch *et al*. (2018), Hasibeder *et al*. (2015), and Schmitt *et al*. (2010).

### Experimental set-up

Four summer drought treatments were established in the enclosed area at the study site: a single year summer drought, three years of recurrent summer drought, 13 years of recurrent summer drought, and ambient conditions (n = 4, respectively). The treatment of 13 years of recurrent drought was established in 2008, the three-year recurrent drought in 2018, and the single-year drought in 2020. Summer drought was simulated for seven weeks from 17 June until 7 August 2020 using rainout shelters (Figure 1). This is the warmest period in the growing season and therefore the period where a natural drought is most likely to occur. The timing and intensity of the annual drought were kept as similar as possible in previous years since 2008. The ambient treatment was not covered with rainout shelters and thus exposed to ambient precipitation. The rainout shelters of the single and three years of recurrent summer drought had a base area of 3 x 3.5 m^2^ and 2.5 m height. Each of these shelters contained a single 20 x 20 cm^2^ plot. The two rainout shelters of the 13 years of recurrent drought were twice the size, resulting in a base area of 3 x 7 m^2^ (shelter height was the same with 2.5 m). These larger shelters contained two 20 x 20 cm^2^ plot, installed at either side of the shelter. In total, 4 plots for ambient, single, three, and 13 years of drought frequency were installed four weeks before summer drought simulation in 2020, respectively. Each plot was located at least 50 cm away from the rainout shelter to avoid edge effects. Rainout shelters with plots were distributed in four blocks in a transect across the mountain meadow.

**Figure 1.**
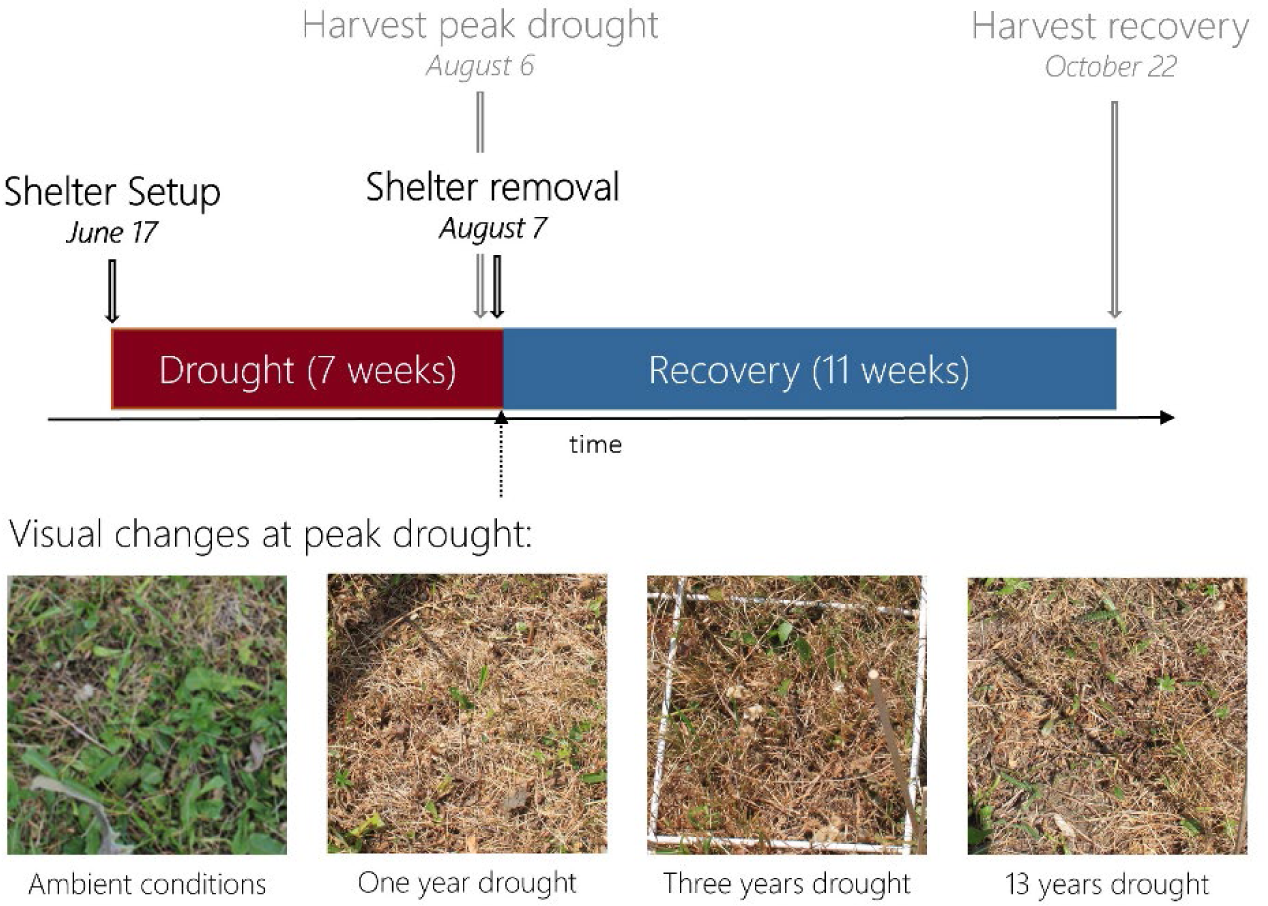
The course of the experiment including setup and removal of the rainout shelters, and peak drought and recovery harvest. The pictures show the plots exposed to ambient conditions, and to one year, three years, and 13 years of drought. All pictures were taken in early August at peak drought.

All shelters were covered with a transparent UV-A and UV-B transmissive plastic foil (Lumisol clear AF, Folitec, Westerburg, Germany, light transmittance ca. 90 %). To allow air circulation, the shelters were open at the bottom (up to 0.5 m) and at the top of the face sides. The drought period was ended by removing the shelters and watering of all 20 x 20 cm^2^ plots with 31 mm of distilled water. At this time point, the entire site was also mown, and the cuttings were removed.

### Measurements

#### Microclimate

A microclimate station at the study site recorded precipitation, air temperature, and relative air humidity (see details in Hasibeder *et al*. (2015)). During the whole season, soil water content (SWC) (Decagon EC-5 and 5TM, Decagon Devices, Pullman, WA, USA; SM300, Delta-T Devices, Cambridge, UK; Hobo S-SMD-M005, Onset Computer corporation, Bourne, MA, USA) was measured continuously in the main rooting horizon (30-min interval) at 5, 10, and 20 cm of soil depth. The measurements were replicated twice in the single-, and three-years treatment, and three to four times in the ambient treatment. Due to partial sensor failures, no data is available for the 13-years treatment. For results on the microclimate and SWC, see Figure S1.

#### Measurements throughout the growing season

In all 16 plots, species composition and vegetation height were measured every two weeks from 22 May to 22 October 2020. Plant species composition was estimated by noting for each species present the number of modules and flowering stems. Modules refer to the number of tillers for grasses and graminoids, the number of rosettes for small dicots, and the number of leaves for larger dicots per rooted species, following (in ’t Zandt *et al*. 2021; Herben *et al*. 2013; Herben *et al*. 1997). Counting was accompanied by measurements of mean vegetation height (in cm). Seedling composition in the plots was counted separately every three weeks between the 23 June and the 22 October 2020. A seedling was defined as a plant individual with less than two leaves, and was not part of the module counts. Seedlings were counted per plant functional group: forbs, grasses, and legumes.

#### Biomass, necromass, and leaf traits

Aboveground biomass (living plant material) and necromass (dead plant material) was sampled destructively in the plots at peak drought on 6 August 2020 and during recovery on 22 October 2020 (Figure 1). As the entire site is mown every summer at the same time point as the peak drought sampling, there was no effect of peak drought biomass removal on recovery sampling. For this, plant material was cut at 2 cm above the soil and frozen at -18 °C until further analysis. Samples were split into plant species and separated into leaves, stems, and flowers. This was done for biomass and necromass, respectively. For each species of each plot, the leaf area was determined from thawed leaves that were saturated with water and scanned (V700 Photo, Epson, WinRHIZO Pro 2012, Regent Instruments). Scanned leaves were dried in the subsequent three days at 60 °C and were weighed. Specific leaf area (SLA) was calculated by dividing the scanned leaf area by the dry weight of the leaves for each species and treatment, respectively. Leaf area index (LAI) was calculated from the leaf surface of all leaves divided by the total unit of ground area of the plot.

We also assessed drought frequency effects on aboveground phytomass fractions. These were calculated as the proportion of biomass, necromass, and flower mass of the total aboveground phytomass in each plot. We did this, because phytomass fractions were in most cases more informative concerning drought frequency effects than absolute values of phytomass (Figure S2) as well as biomass and necromass (data not shown). This is because, a substantial portion of the phytomass at the subalpine meadow is produced early in the season between May and mid-July (data not shown), which is before the peak drought period. Hence, in our study, (recurrent) drought had little effect on this early-seasonal phytomass build-up. Instead, increasing drought frequency resulted in phytomass loss and/or plant mortality, which is visible in *fractions* of the phytomass, as it illustrates the redistribution of mass. Notably, while biomass refers to productivity, biomass fraction reflects the maintained productivity.

#### Plant species seasonality and flowering

We assessed plant species seasonal growth to characterise plant species as early and late seasonal species. For this, we determined the time point at which the plant species peaked in its number of modules. The typical seasonal behaviour of each plant species was determined based on the ambient treatment plots only. Early seasonal species were defined as species that peaked in modules before the biomass cut and thus reached peak growth between the end of May to end of July. Late seasonal species were defined as species that peaked in modules after the biomass cut and thus reached peak growth between August and October. Shifts in early and late seasonal patterns of species were assessed using values from all treatments.

Because of the low occurrence of flowers, seasonal shifts in flowering was calculated based on flower biomass. For this, we calculated the ln-response ratio between a species’ flower dry mass at peak drought and its flower dry mass after the 11-week recovery period. A positive value signifies plant species flowering early in the growing season, i.e., during the drought period. A negative value signifies plant species flowering late in the growing season, i.e., during the recovery phase.

#### Plant community metrics and composition

Plant community evenness throughout the season was calculated as Pielou’s index using the plant species module counts per time point and plot. For this, we divided the plant diversity Shannon index by the ln-transformed species richness using the function diversity and specnumber of the vegan package (R Core Team 2021; Oksanen *et al*. 2020). Plant community seasonal synchrony was calculated following Loreau & De Mazancourt (2008), using the package codyn (Hallett *et al*. 2016). In brief, we compared the variance of the total module counts of all species over the season (𝑥_𝑇_) with the summed variances of the individual species’ module counts over the season (𝑥_𝑖_) to quantify seasonal synchrony:

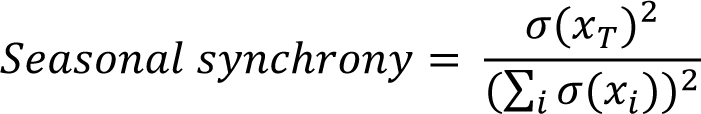

Where:

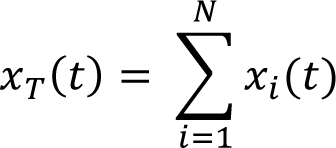

With 𝑥_𝑇_ (𝑡) being the total module counts of all species combined at time *t*, 𝑥_𝑖_ (𝑡) the module counts of species *i* at time *t*, 𝜎(𝑥_𝑇_) ^2^the variance of the total module counts of all species over the season combined, and 𝜎(𝑥_𝑖_) the variance of the module counts of species *i* over the season. Using the codyn package (Hallett *et al*. 2016), plant species loss between subsequent time points was calculated to indicate species loss *within* the growing season. To determine whether communities were dominated by either early or late plant species, we calculated the proportional number of early plant species present in each community at each time point (see section Plant species seasonality and flowering).

We assessed plant community composition using Non-Metric Multidimensional Scaling (NMDS) using metaMDS from the vegan package (Oksanen *et al*. 2020). NMDS on both module counts and biomass were run on Bray-Curtis dissimilarity matrices between plots and time points and with four dimensions to minimise stress values.

#### Statistics

All calculations and statistical analyses were done in R 4.1.1 (R Core Team 2021). Effects of drought frequency on percentage data (phytomass fractions) were performed using Generalized Linear Mixed Models (GLMM) with a beta distribution and logit-link using the package glmmTMB (Magnusson *et al*. 2017). Drought frequency effects on species richness and the number of seedlings were tested using Generalised Linear Mixed-Effects Models (GLMER) with a poisson distribution and log-link function using the lme4 package (Bates *et al*. 2015). Drought frequency effects on continuous variables (seasonal synchrony and community evenness) were tested using Linear Mixed Effects Models (LMER) using the lme4 package (Bates *et al*. 2015). Models including ‘date’ (continuous, scaled) were run with ‘plot’ as a random effect to take the repeated sampling design into account. Models on plant species-specific traits were only run for plant species for which at least two data points occurred in each treatment.

Heterogeneity of the mountain meadow was taken into account as a random effect in statistical models. For models on plant community measures (phytomass fractions, seasonal synchrony, species richness, community evenness and seedling numbers), ‘block’ was added as a random effect. Models on plant species-specific traits included the mean vegetation height of each plot before the setup of the rainout shelters as a random effect. This variable was better at capturing variation *within* drought treatments than ‘block’ and, importantly, was not significantly affected by previous drought events (data not shown).

For GLMM and GLMER models, we tested over- and under-dispersion of the model residuals using the package DHARMa (Hartig 2022). For LMER models, we tested assumptions of homogeneity of variances, and normal residual distributions following Zuur *et al*. (2010). Data was ln- or sqrt-transformed when model residuals did not adhere to model assumptions.

For plant community composition based on NMDS analyses, significant separation of communities resulting from ‘drought frequency’ (factor), date (scaled, continuous), and their interaction was tested using a restricted permutation test: samples were permuted *within* plots to take the repeated sampling structure over time into account. The test was performed using adonis from the vegan package with 999 permutations (Oksanen *et al*. 2020). A passive overlay of ‘drought frequency’ (ln-transformed + 1), ‘species richness’, ‘proportion early species’, ‘seasonal synchrony’, species loss between subsequent time points (for NMDS based on counts only), ‘LAI’, functional group total abundance, flower abundance, and seedling abundance was created and tested for significance using the function envfit of the vegan package with 999 permutations (Oksanen *et al*. 2020).

## Results

### Drought frequency decreased biomass and species richness and increased necromass

In a species-rich sub-alpine mountain meadow, we asked how an increase in the annual frequency of severe summer drought affects plant community phytomass, species richness, and flower production during peak drought and recovery. At the peak of drought, 7 weeks after the setup of the rainout shelters, we found that drought frequency significantly decreased the fraction of community biomass and increased the fraction of community necromass. These effects were non-linear, with the strongest effects observed during low drought frequency, plateauing as drought frequency increased (Figure 2AB). The increase in necromass was primarily related to an increase in the necromass fraction of grass and legume species (Figure S3). Eleven weeks after soil rewetting (recovery), these effects were still present, albeit to a lesser extent (Figure S4AB). In the recovery phase, the fraction of forb biomass decreased, while the fraction of grass biomass and necromass increased with drought frequency (Figure S3).

**Figure 2.**
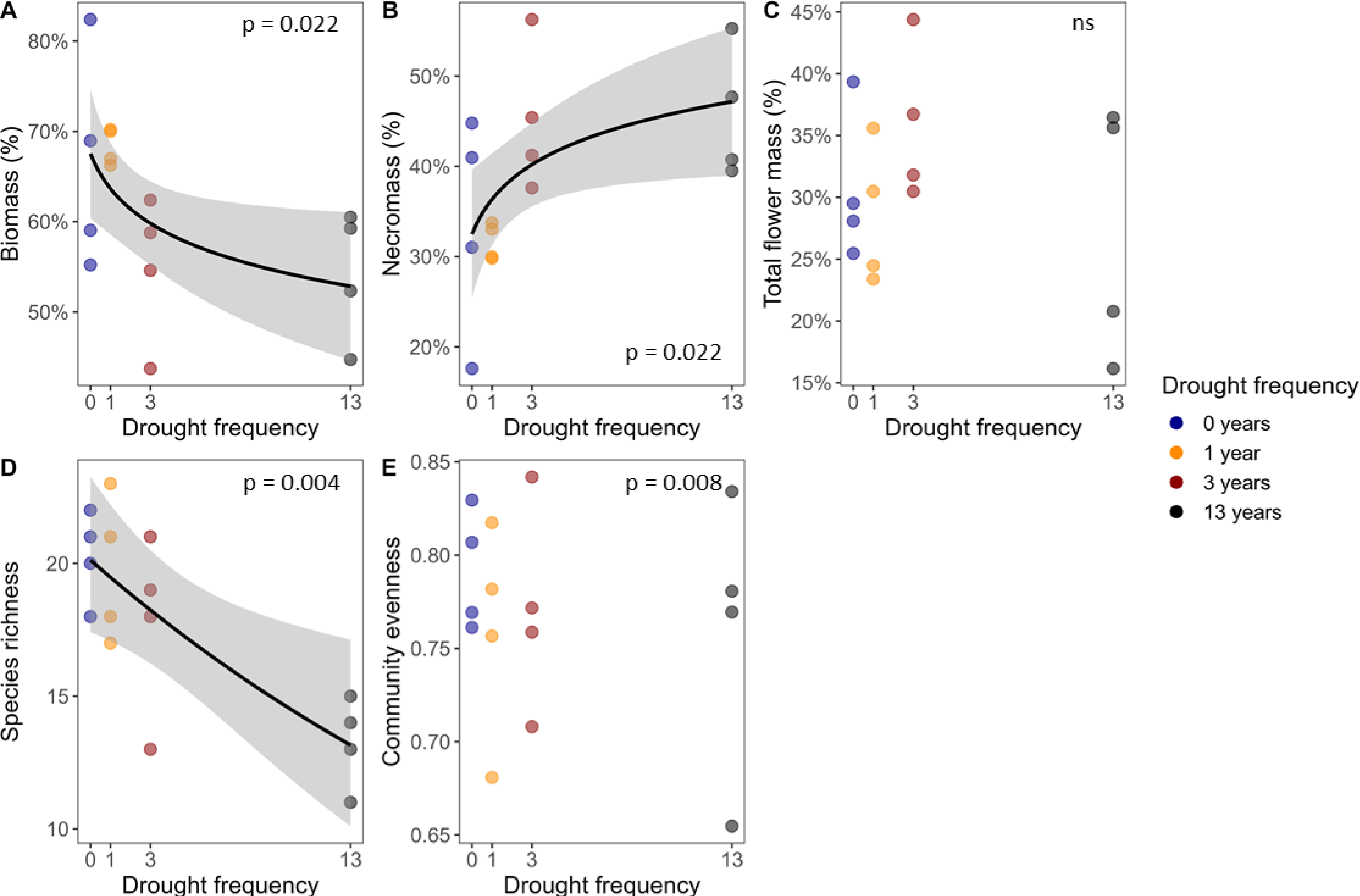
Drought frequency effects at peak drought on (A) the fraction of plant community biomass (living plant material), (B) the fraction of plant community necromass (dead plant material), (C) the fraction of total flower mass, (D) plant community species richness, and (E) plant community evenness. A-C are expressed as percentages of the phytomass (bio- and necromass) in each plot. For significant relations, the black line indicates the mean relation with the shading showing the 95% confidence interval. P-values of significant relations are shown (ns: no significant effect, p > 0.05, n = 4 per drought frequency level). For effects of drought frequency on overall plant community parameters in the recovery phase, see Figure S4.

Drought frequency did not significantly affect the fraction of flower biomass at peak drought (Figure 2C). During recovery, however, the fraction of flower biomass was significantly decreased with increasing drought frequency (Figure S4C).

Furthermore, drought frequency significantly decreased plant species richness at both peak drought and recovery (Figure 2D; S4D). On average, 13 years of recurrent summer drought resulted in a 35 % and 46 % loss of plant species compared with ambient conditions at peak drought and recovery, respectively. Despite this loss in plant species, drought frequency did not significantly affect plant community evenness at both peak drought and recovery (Figure 2E; S4E).

### Drought frequency decreased forb and legume seedlings

To assess the impact of drought frequency on the establishment of plant seedlings throughout the growing season, we monitored the number of seedlings per plant functional group at biweekly intervals throughout the season. Drought frequency affected the number of forb and legume seedlings, but not of grass seedlings. During drought and recovery, the number of forb and legume seedlings was consistently lowest with 13 years of recurrent summer drought (Figure 3). Additionally, while in the ambient treatment and one-year drought plots, many forb and legume seedlings appeared after the mowing event, these peaks were lower in plots with three years and especially 13 years of recurrent summer drought (Figure 3). At the same time, plant community LAI decreased with drought frequency at both peak drought and recovery (Figure S5AB). The latter indicates that the soil surface was less covered by leaves of the established plant community with an increase in drought frequency.

**Figure 3.**
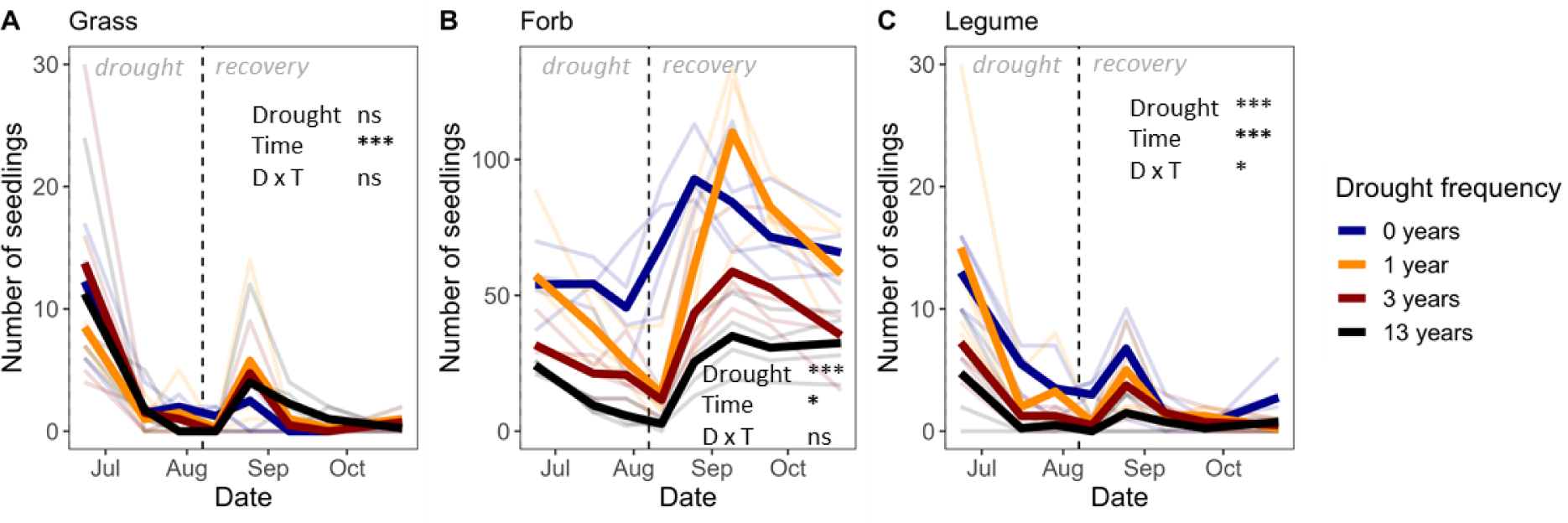
Drought frequency effects on the number of seedlings of (A) grass, (B) forb, and (C) legume species throughout the growing season. The dotted line indicates the timepoint at which drought ended, and the plots were mown and rewetted. Solid lines indicate mean seedling numbers, and faded lines indicate seedling numbers of each plot (n = 4 per drought frequency treatment). Results from analyses of variances are shown with ‘***’ indicating p < 0.001, ‘*’ indicating 0.01 > p < 0.05, and ‘ns’ indicating no significant effect (p > 0.05).

### Drought frequency increased plant community seasonal synchrony

We tested how drought frequency affected plant community seasonal dynamics based on plant species module counts which we monitored throughout the growing season. The total number of plant modules in all treatments fluctuated throughout the growing season (Figure S6A). However, it was not significantly affected by drought frequency throughout the season (Figure S6A). Furthermore, plant species richness before, during, and after drought was consistently lowest in plots subjected to 13 years of drought, followed by three years, and one year of drought (Figure S6B). Plant species richness across all treatments dropped sharply after peak drought and the mowing event but increased to levels similar to their respective pre-drought conditions within a month of recovery (Figure S6B).

Next, we tested drought frequency effects on plant community seasonal synchrony, which we calculated based on the module counts of each plant species over the season. We found that drought significantly increased seasonal synchrony, that is, drought synchronised the timing of plant species growth during the season. Increasing drought frequency enhanced this synchronising effect. This relationship was non-linear, with the strongest increase in synchrony occurring during low drought frequencies and the increase plateauing as drought frequency became higher (Figure 4A). Furthermore, seasonal synchrony was negatively related to the average plant species richness throughout the season, that is, synchrony was high when average plant species richness was low (Figure 4B).

**Figure 4.**
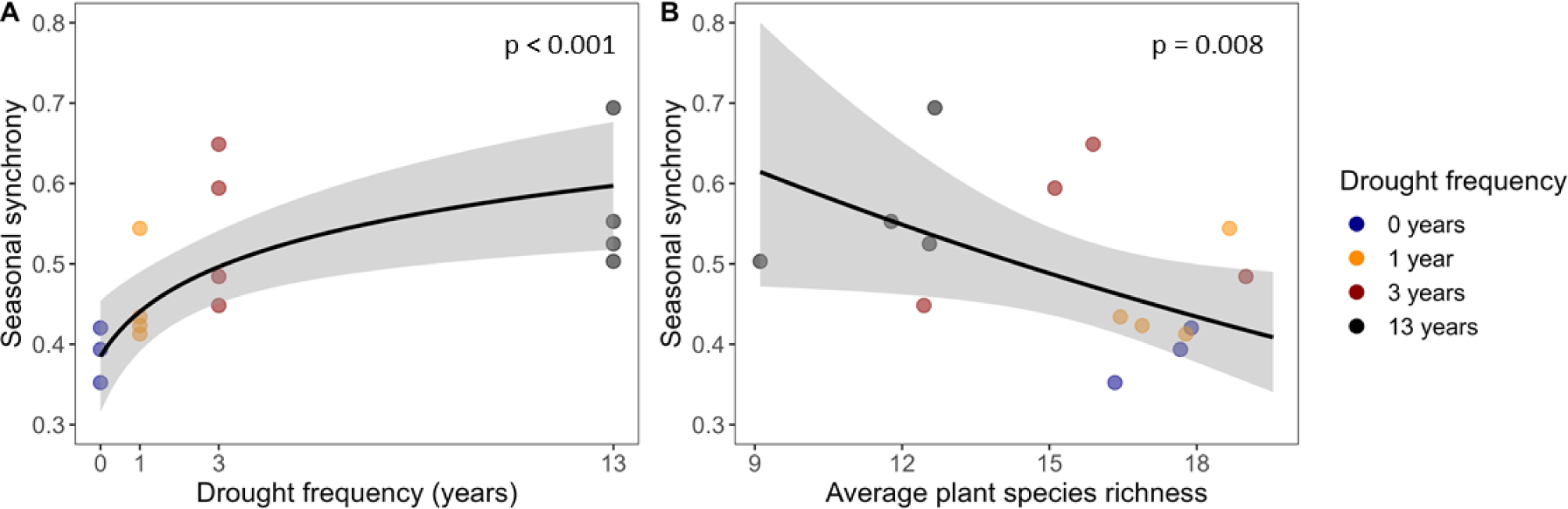
(A) Effects of increasing drought frequency on seasonal synchrony (sensu Loreau & De Mazancourt (2008)) based on plant module counts across the growing season from mid-May to mid-October, and (B) seasonal synchrony in relation to the average plant species richness (n = 4 per drought frequency treatment). The black line indicates the mean relation with the shading showing the 95% confidence interval. P-values of significant relations are shown.

### Drought frequency shifts plant community composition towards less early seasonal and more grass species

We examined the effect of drought frequency on plant community composition throughout the season using non-metric multidimensional scaling (NMDS). We found significant differentiation of plant communities based on drought frequency, with communities experiencing no drought and 13 years of drought exhibiting the greatest dissimilarity (Figure 5A, S7). This compositional separation, driven by drought frequency, was primarily captured along the first axes of the NMDS plot. This first axis coincided with the decrease in plant species richness and the increase in community synchrony as drought frequency increased (Figure 5A). Moreover, we observed that the compositional shifts associated with increasing drought frequency were characterised by a high grass abundance and a decrease in early-seasonal plant species. In addition, LAI, the number of forb flowers, and the number of forb seedlings decreased with drought frequency (Figure 5A). These patterns were similar when basing the analyses on plant biomass collected at peak drought and recovery (Figure S8).

**Figure 5.**
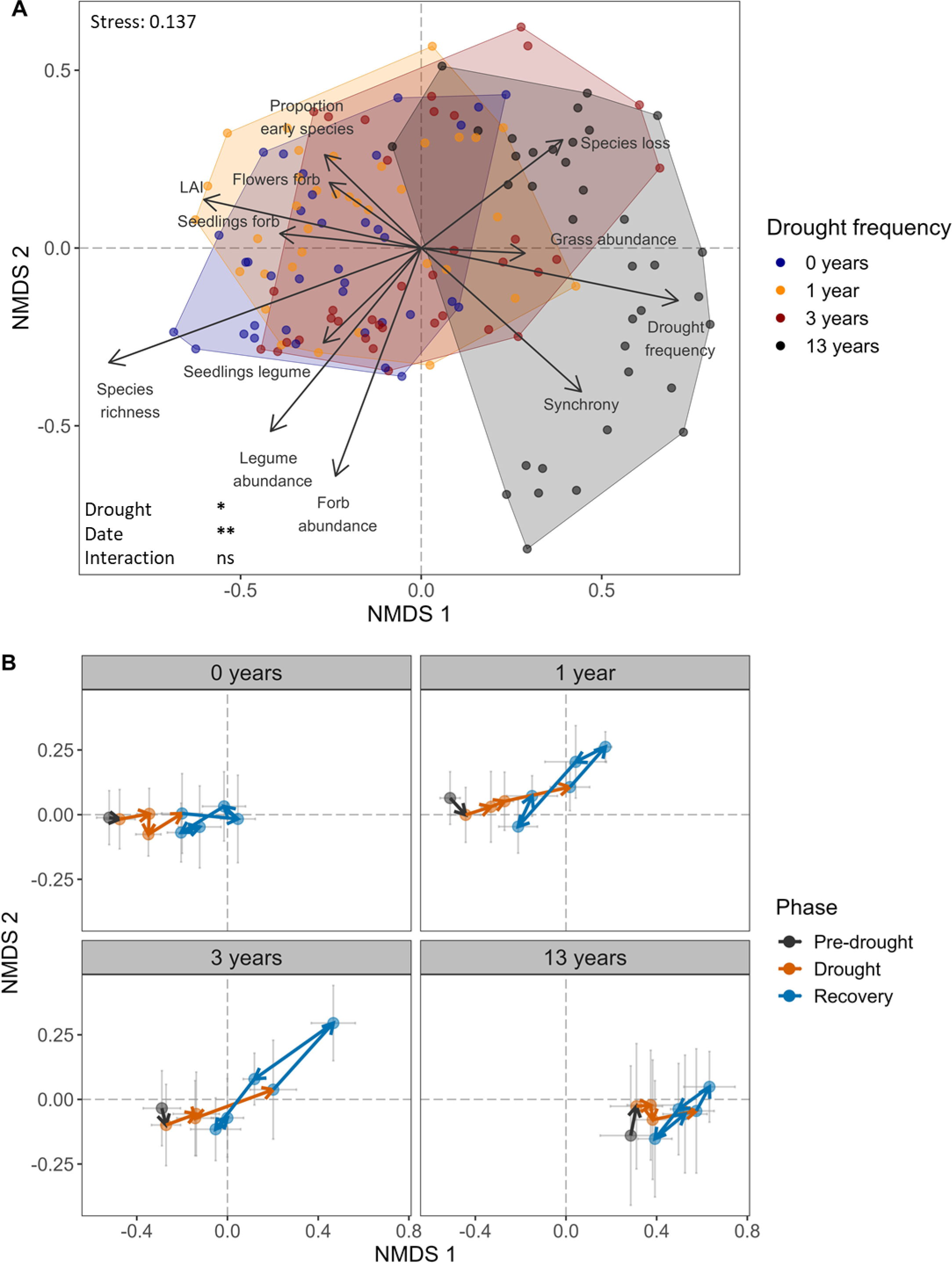
Non-metric multidimensional scaling (NMDS) of plant species module counts throughout the growing season in permanent plots receiving no drought (0 years), one year, three years, and 13 years of recurrent summer drought. (A) Drought frequency effects on plant community composition with a passive overlay of community variables significantly relating to the NMDS patterns. Each point represents a plot at a point in time. Hulls group all plots and time points per drought frequency treatment together. Length of arrows indicate the strength of the relation. ‘Proportion early species’ refers to the proportion of early species versus the total number of species present in a plot. ‘Grass’, ‘forb’, and ‘legume abundance’ refer to the total number of modules summed per functional group. ‘Synchrony’ refers to community seasonal synchrony sensu Loreau & De Mazancourt (2008). ‘Species loss’ refers to the percentage of species lost within the growing season between two subsequent time points (species gain was not significantly related). ‘LAI’ refers to leaf area index. Results of a restricted permutation test is shown with ‘**’ indicating 0.001 > p < 0.01, ‘*’ indicating 0.01 > p < 0.05, and ‘ns’ indicating no significant effect (p > 0.05) (n = 4 plots per drought frequency treatment). (B) Community compositional development over time. Points are mean values ± standard errors. Note that axes scales differ between A and B to improve readability. For NMDS axes 3 and 4, see Figure S7. For position of plant species along the axes, see Figure S9. For the plant community composition results based on plant species biomass, see Fig S8.

During the recovery phase, plant communities in the plots exposed to one and three years of drought shifted along the second NMDS axis. After rewetting, these one- and three-year droughted plots temporarily shifted into the top-right quadrant of the NMDS (Figure 5B). This direction was related to a high loss of species, particularly early-seasonal species (for specification see Table S1), and a decrease in forb and legume abundance (Figure 5B). However, later in the recovery phase, the one- and three-year droughted communities gradually returned to a similar position on the second axis of the NMDS plot as before the recovery phase (Figure 5B). This recovery pattern indicates slower recovery of early-seasonal forb, and legume species within the one- and three-year droughted plots (Figure 5B).

### Weak drought frequency effects on individual plant species responses

Finally, for 10 plant species that were present in sufficient abundances in all three drought frequency treatments and the ambient treatment, we tested whether drought frequency affected SLA at peak drought and recovery, as well as the timing of their growth in the season. At peak drought, we found that drought frequency significantly decreased SLA of *Alchemilla vulgaris, Leontodon hispidus, Plantago lanceolata* (marginally)*, Trifolium repens,* and *Veronica chamaedrys* (Figure S10). During recovery, these patterns had disappeared (Figure S11).

Drought frequency only significantly affected the timing of *Trifolium pratense,* which peaked in number of modules earlier in the season with a high drought frequency (Figure S12). Drought frequency only significantly affected the timing of flowering of *Leontodon hispidus,* which shifted its flower production to earlier in the season with a high drought frequency (Figure S13).

## Discussion

We investigated the effects of recurrent summer drought of up to 13 years on plant communities in a sub-alpine mountain meadow, as well as their underlying mechanisms. We found that drought frequency increased plant community seasonal synchrony. This increase in synchrony with greater drought frequency was underpinned by a decrease in species richness, a loss of early-seasonal plant species, a shift in plant functional groups, and the constrained establishment of seedlings throughout the growing season. Moreover, these changes were associated with a decreased biomass fraction as drought frequency increased.

### Higher seasonal synchrony with increasing drought frequency due to species loss, compositional shifts, and lower seedling establishment

We found that with increasing drought frequency, species **synchrony** within the plant community was enhanced in a non-linear fashion: the increase was strongest at low drought frequency and plateaued at high drought frequencies. This indicates that the species in the plant community under increasing drought frequency were more similar in their timing of plant species growth throughout the season. Results on increasing synchrony *between* years for grasslands were reported by other studies, finding an increase in synchrony with drought frequency of three years in an alpine meadow (He *et al*. 2022), and four years across grasslands (Muraina *et al*. 2021). Our study suggests that with increasing drought frequency (to up to 13 years of recurrent drought) seasonal synchrony becomes increasingly pronounced. In consequence, multiple droughts may hamper the stability (De Mazancourt *et al*. 2013; Loreau & De Mazancourt 2013), e.g. under future disturbances. This is because synchronous communities are typically less stable to disturbances than asynchronous communities (Luo *et al*. 2023b; Muraina *et al*. 2021; Craven *et al*. 2018; De Mazancourt *et al*. 2013). Hence, an increase in seasonal synchrony with drought frequency suggests that multiple droughts may hamper the stability of communities to future disturbances. We suggest that with the ongoing and projected increases in the frequency of drought events due to climate change (IPCC 2021), grassland ecosystems become increasingly seasonally synchronous and vulnerable to future disturbances.

We identified several underlying mechanisms that contributed to the increase in synchrony with increasing drought frequency. First, the enhancement in synchrony was associated with a **decrease in plant species richness**. This drought-induced species loss aligns with findings from other studies that observed enhanced mortality during and following single (Sippel *et al*. 2018; Stampfli *et al*. 2018; Frank *et al*. 2015), and recurrent drought events (Luo *et al*. 2023a; Wagg *et al*. 2017). Species richness mostly tends to be negatively associated with synchrony (Zhang *et al*. 2018; Hautier *et al*. 2014), but was also found to be positively associated with synchrony (Valencia *et al*. 2020). The negative association between plant species richness and seasonal synchrony in our study likely occurred because the plant species that survived in the community with recurrent drought, were more similar in terms of their seasonal development. Moreover, under increasing drought frequency, communities were dominated by species growing later in the season, as **early-growing species were lost**. This may have been caused by a reduced recovery potential of early-season species after the drought event: We found that early-seasonal species were taking a longer time to recover under single- and three-year drought. While it was suggested, that early-season species may be more susceptible to drought (Mamolos *et al*. 2001), we found first evidence that early-growing species actually get lost at high drought frequencies. As a result, under recurrent drought, more temporally similar growing species lead to more seasonally synchronized communities.

Furthermore, the seasonal synchronization of species may have additionally been driven by the observed shift within plant **functional groups** under higher drought frequency. We found that the community composition showed a reduction in forb and legume species, and more grass species with increasing drought frequency. The loss of species within these functional groups with increasing drought frequency may be explained by the observed lower forb and legume seedlings throughout the season and the slower recovery of forbs and legumes post drought. Hence, our findings support the notion that forb and legume species are less robust to recurrent drought compared with grass species. This is in line with studies on single drought (De Boeck *et al*. 2018; Hoover *et al*. 2014) and recurrent drought events (Luo *et al*. 2023a; Luo *et al*. 2023b; Sun *et al*. 2023; Xu *et al*. 2021), which also suggest grass species to be less sensitive to drought compared to forbs. Overall, the observed shifts under recurrent drought towards less diversity in functional groups can enhance synchrony by increasing the similarity of seasonal growth within the community. This similarity in functional groups can furthermore decrease the variety in responses to drought throughout the growing season (Zhang *et al*. 2020) and thereby contribute to an enhanced vulnerability under future disturbances.

The increase in synchrony with higher drought frequency may also be driven by the **constrained establishment of seedlings** throughout the growing season: As drought frequency increased, we observed a **decline in seedling numbers**. This occurred during the entire vegetation period and was most pronounced with high drought frequency of 13 years. At the same time, less soil surface area was covered by leaves under high drought frequency. Hence, the seedling decline took place despite a diminished light limitation, and thus likely a higher chance of seedling survival in the 13-year recurrent drought plots (Eskelinen *et al*. 2022). Moreover, we observed that plant species **reproductive output** was inhibited with increasing drought frequency: In the recovery phase, the fraction of flower biomass was decreased with increasing drought frequency. A negative drought effect on reproductive output was also found by Zeiter *et al*. (2016). Reasons might be, that species experiencing drought were unable to produce flowers due to preceding and ongoing water limitation. Consequently, in the communities with these high drought frequencies, fewer seedlings appeared in the recovery phase.

Overall, in our study, flower development and consequently seedlings, were severely inhibited with higher drought frequencies. This reduces the establishment of seedlings and leads to species loss, with effects on community composition and synchrony. During long-term recurrent drought, effects of drought on reduced plant reproductive output and resultant seedlings likely exacerbated plant species loss and the increase in seasonal synchrony. However, single- and low-frequency experiments do not cover drought effects that go beyond the lifetime of the plant species. We therefore propose that, to comprehensively comprehend changes in plant communities arising from high drought frequency, the implementation of long-term experiments is essential.

### Enhanced loss of biomass fraction with increasing drought frequency

Drought did not reduce productivity in our study (Figure S2), which is consistent with findings on productivity under five years of drought by Jentsch *et al*. (2011) and shoot biomass under four years of drought by Fuchslueger *et al*. (2016). Nonetheless, other low drought frequency studies reported a continuous decline in aboveground net primary productivity under recurrent drought of up to four years (Luo *et al*. 2023a; Xu *et al*. 2021). The divergent results may be attributed to the different frequencies of drought events or plant communities. However, we found that increasing drought frequency gradually decreased the fraction of plant community biomass and enhanced the fraction of necromass at peak drought and recovery, respectively. These were non-linear effects, with strongest increases/decreases at low drought frequency that plateaued as drought frequency became higher. Hence, during the growing season, including drought and recovery, the maintained productivity is lower, and senescence is higher with increasing drought frequency. The higher senescence at peak drought and recovery supports our observation of increasing species loss throughout the growing season. Reductions in maintained productivity might be caused by the lasting repercussions of drought on plant community composition and synchrony.

Under recurrent drought, plants have the capacity to develop **adaptations** that can dampen drought effects (Müller & Bahn 2022). This can occur e.g., by adjusting photosynthesis, osmosis (Li *et al*. 2022), root biomass (Legay *et al*. 2018), epigenetics, transcription factors and rate (Jacques *et al*. 2021; Ding *et al*. 2012; Bruce *et al*. 2007), as well as transgenerational changes (Chen *et al*. 2022). Within our study, we found only weak species adaptations under recurrent drought. For example, increasing drought frequency did not result in adaptations of the duration of the growing season and only shifted the timing of growth for one legume species to earlier in the season. Also, the timing of plant species flowering was only shifted earlier with increasing drought frequency for one forb species. Moreover, the lower SLA at peak drought of some of the most abundant forb species and one legume species with increasing drought frequency can hint towards higher water-use efficiency, resource conservation, and reduced plant growth, which can enhance short-term stress tolerance (Wellstein *et al*. 2017). This short-term adaptation could lead to less vulnerability under drought. Overall, adaptations did not play a major role in compensating drought effects in our study.

Chen *et al*. (2022) suggested that adaptations *along generations* under recurrent drought lead to improved ecosystem responses. However, we suggest that the lower reproduction might overrule any potential adaptive effects of single plant species along generations through species loss. Plant communities are therefore likely not able to adapt to the predicted rapid increase in drought frequency with the ongoing change in climate. To gain further understanding of the repercussions on plant community due to increasing drought frequencies, long-term drought experiments, including transgenerational effects, could yield important insights.

## Conclusion

We show that long-term annual drought frequency in grasslands increases plant community seasonal synchrony. Synchrony increased through a reduction in species richness, a loss of early-seasonal plant species, a shift in plant functional groups, and the constrained establishment of seedlings throughout the growing season. These changes were associated with a decreased fraction of biomass with increasing drought frequency.

Overall, we conclude that with the increase in frequency and severity of drought events due to climate change, grassland plant communities will increase in seasonal synchrony. As asynchrony is a critical component of community stability, we suggest that this will lead to lasting effects far beyond the drought event, by increasing the vulnerability of ecosystems to future disturbances. Moreover, we show that negative drought effects were enhanced with an increasing drought frequency, though effects may vary across ecosystems and climate conditions (Tielbörger *et al*. 2014). We conclude that single and low-frequency drought studies may not adequately predict longer-term changes in our rapidly shifting climate. Furthermore, on the long-term, negative effects, mostly associated with species loss, will likely overrule the comparatively weak adaptation of species. Hence, with the current speed of climate change, communities will be severely and probably irrevocably affected the moment species are lost from the ecosystem. We therefore conclude that plant communities are unlikely to be able to adapt to recurrent drought. Importantly, long-term experiments will be essential in understanding the severity of these climate change impacts.

### Sample CRediT author statement

**Lena M. Müller**: Conceptualization, Investigation, Project administration, Writing - Original Draft (lead), Visualization, Funding acquisition; **Michael Bahn**: Conceptualization, Resources, Funding acquisition, Writing - Review & Editing; **Maximillian Weidle**: Investigation, Writing - Review & Editing; **Georg Leitinger**: Writing - Review & Editing, Funding acquisition; **Dina in ‘t Zandt**: Conceptualization, Investigation, Formal Analysis, Data Curation, Visualization, Writing - Review & Editing, Writing - Original Draft, Supervision

## Acknowledgements

We are grateful to numerous colleagues, students, and friends who have helped to maintain the long-term experiment over the years, as well as to those that assisted in setting up and conducting the sampling for the experiment presented in this study. Amongst others, we thank Beverley Anderson, Mario Deutschmann, Andrew Giunta, Roland Hasibeder, Johannes Ingrisch, Florian Oberleitner, Natalie Oram, David Reinthaler, Michael Schmitt, Carmen Telser, and Herbert Wachter.

The study site is part of the LTSER platform Tyrolean Alps, belonging to the national and international long-term ecological research networks (LTER-Austria, LTER Europe and ILTER). The work was funded by the Forschungszentrum Berglandwirtschaft (project no. 326572). The long-term drought treatment was maintained by projects funded by the Austrian Science Fund (FWF, project no. P22214-B17, P31132, and I1056) and the Austrian Academy of Sciences (ESS-programme, project ClimLUC).

LM was supported by the University Innsbruck through a doctoral scholarship (Doktoratsstipendium aus der Nachwuchsförderung der Universität Innsbruck) and partially funded by the Austrian Science Fund (FWF): [I4969-B]. DZ was supported by a PPLZ Postdoctoral Fellowship awarded by the Czech Academy of Sciences (RVO 67985939).

## Data Availability Statement

Upon acceptance, all data and code (R scripts) will be made publicly available via the Zenodo data repository, will be referenced using a stable object identifier (DOI) in the Materials and Methods section, and cited in the reference list.

