## Supplementary material for "Recurrent drought increases grassland community seasonal synchrony"

### Supplementary Figures and Table

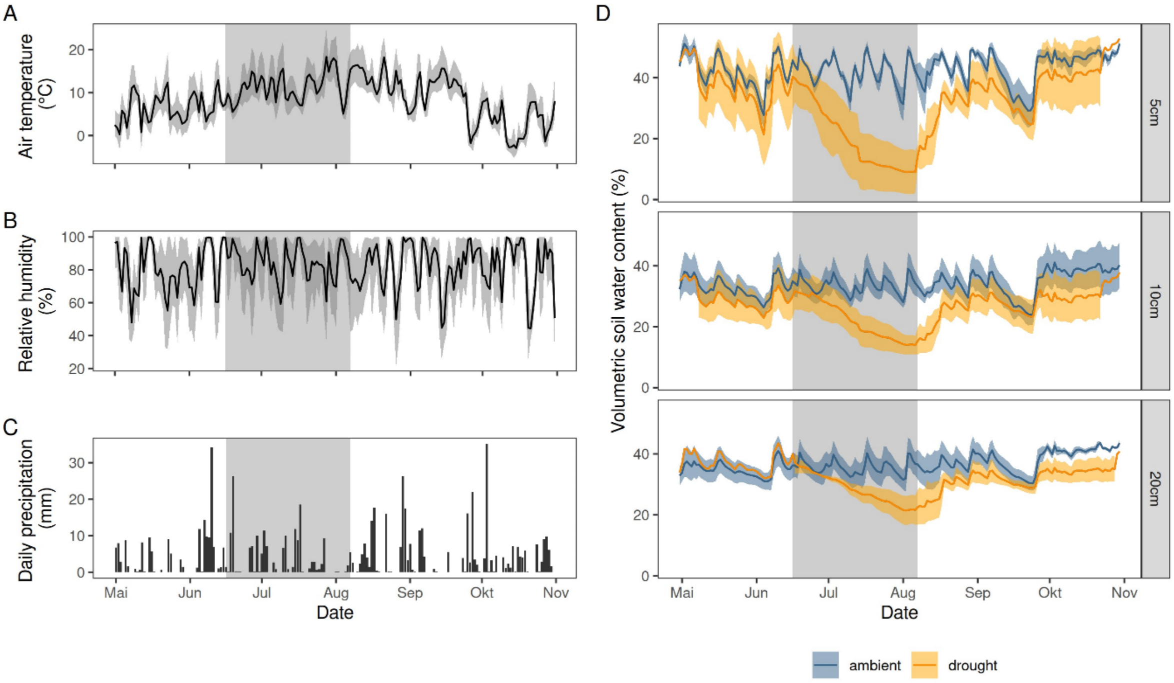


**Fig S1** Microclimate: (A) daily mean air temperature, (B) daily mean relative air humidity with the ribbons indicating the daily range (minimum to maximum values), respectively, and (C) the daily sums of precipitation. (D) Volumetric soil water content (SWC) across treatments in 5, 10, and 20 cm depths, from 1st May to 31st of October, respectively. SWC data is presented for ambient (n = 3-4 per depth), and drought treatment (one and three years of drought grouped; n = 3-4 per depth). Due to partial sensor failures, not all sensor data is available. Individual sensors are summarized to daily means. For each depth and treatment (control and drought), lines indicate means, and ribbons standard deviation, respectively. As soil sensors were not rewetted after peak drought, SWC of drought plots increase along with natural precipitation post-drought.

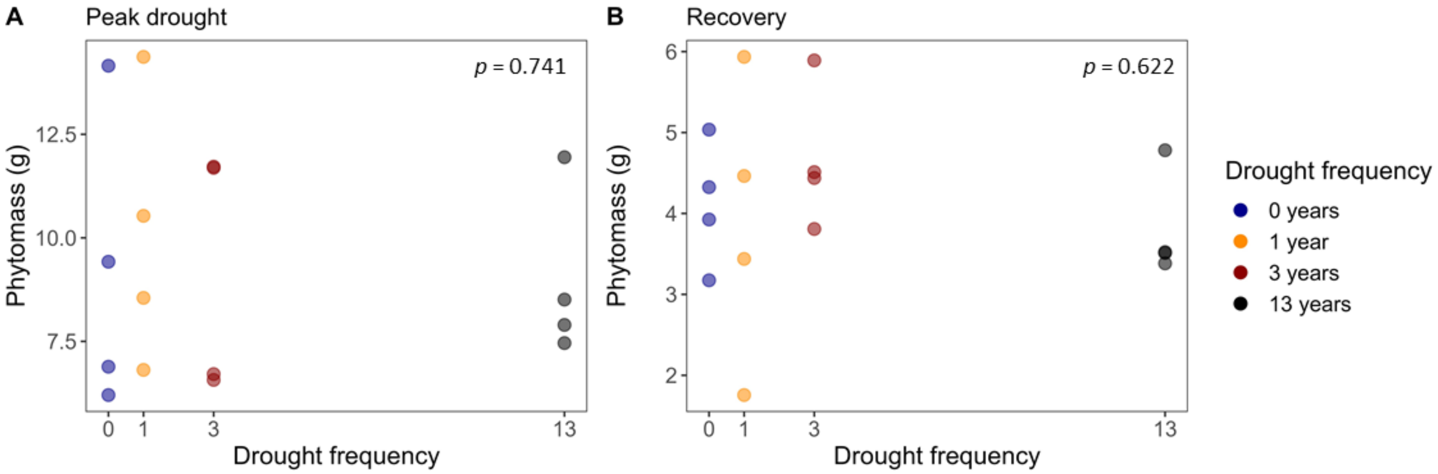


**Fig S2** Drought frequency effects on phytomass (bio- and necromass) at (A) peak drought, and (B) recovery. P-values indicate results from linear mixed effect models (n = 4 per treatment).

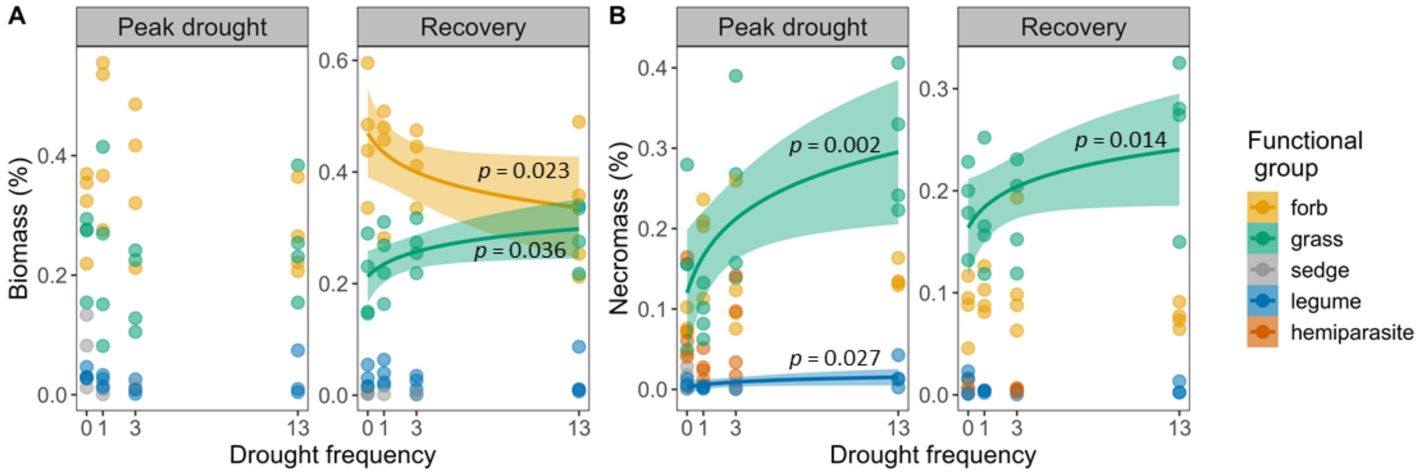


**Fig S3** Drought frequency effects on (A) the fraction of biomass (alive plant material) and (B) the fraction of necromass (dead plant material) of functional plant groups at the peak drought and recovery harvest time points. Fractions are expressed as a percentage of the phytomass (bio- and necromass) per plot. For significant relations, the mean relation with 95% confidence intervals are shown (n = 10-16 per functional group). No alive material of hemiparasites occurred at either time points.

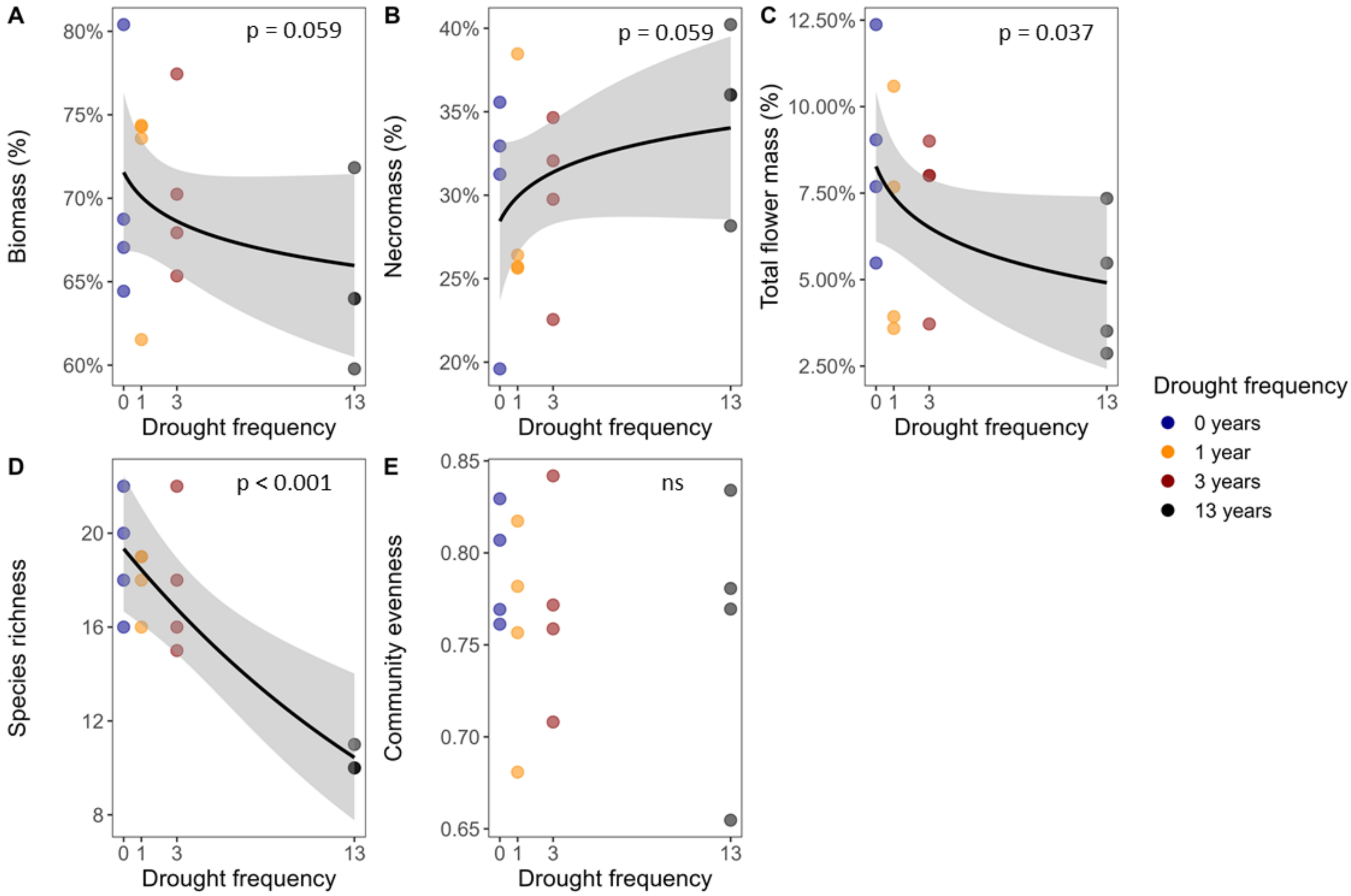


**Fig S4** Drought frequency effects at recovery (11 weeks post drought) on (A) the fraction of plant community biomass (living plant material), (B) the fraction of plant community necromass (dead plant material), (C) the fraction of total flower mass, (D) plant community species richness, and (E) plant community evenness. A-C are expressed as percentages of the phytomass (bio- and necromass) in each plot. For significant relations, the black line indicates the mean relation with the shading showing the 95% confidence interval. P-values of significant relations are shown (ns: no significant effect, $p > 0.05$, $n = 4$ per drought frequency level.

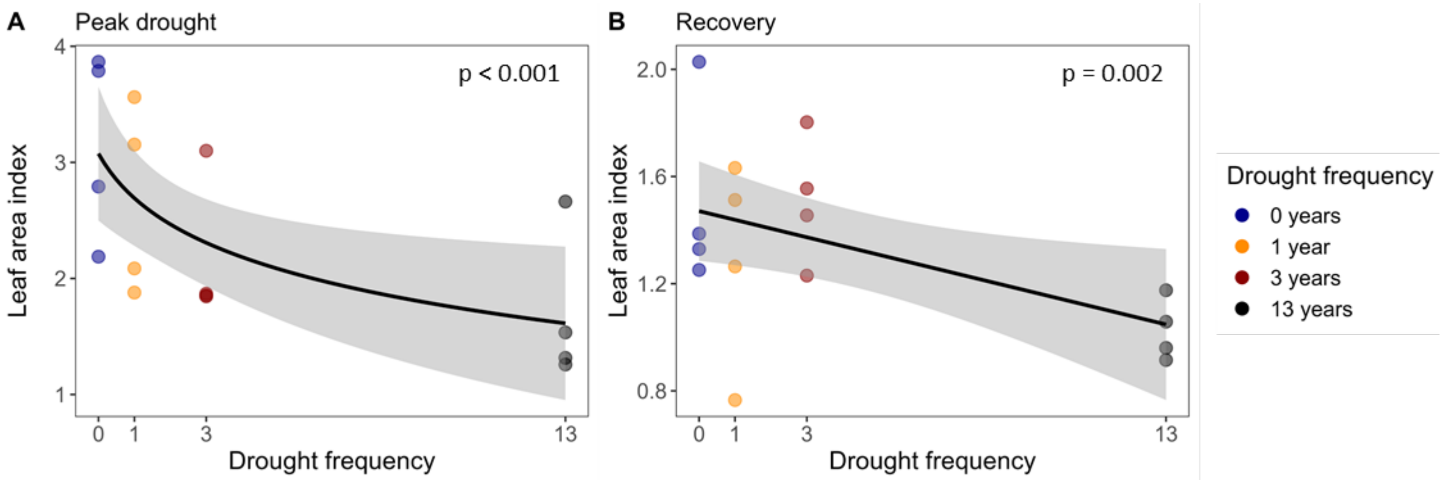


**Fig S5** Drought frequency effects on plant community leaf area index (LAI) during (A) peak drought, and (B) recovery (11 weeks post drought). The black line indicates the mean relation with the shading showing the 95% confidence interval. P-values of significant relations are shown (n = 4 per drought frequency level).

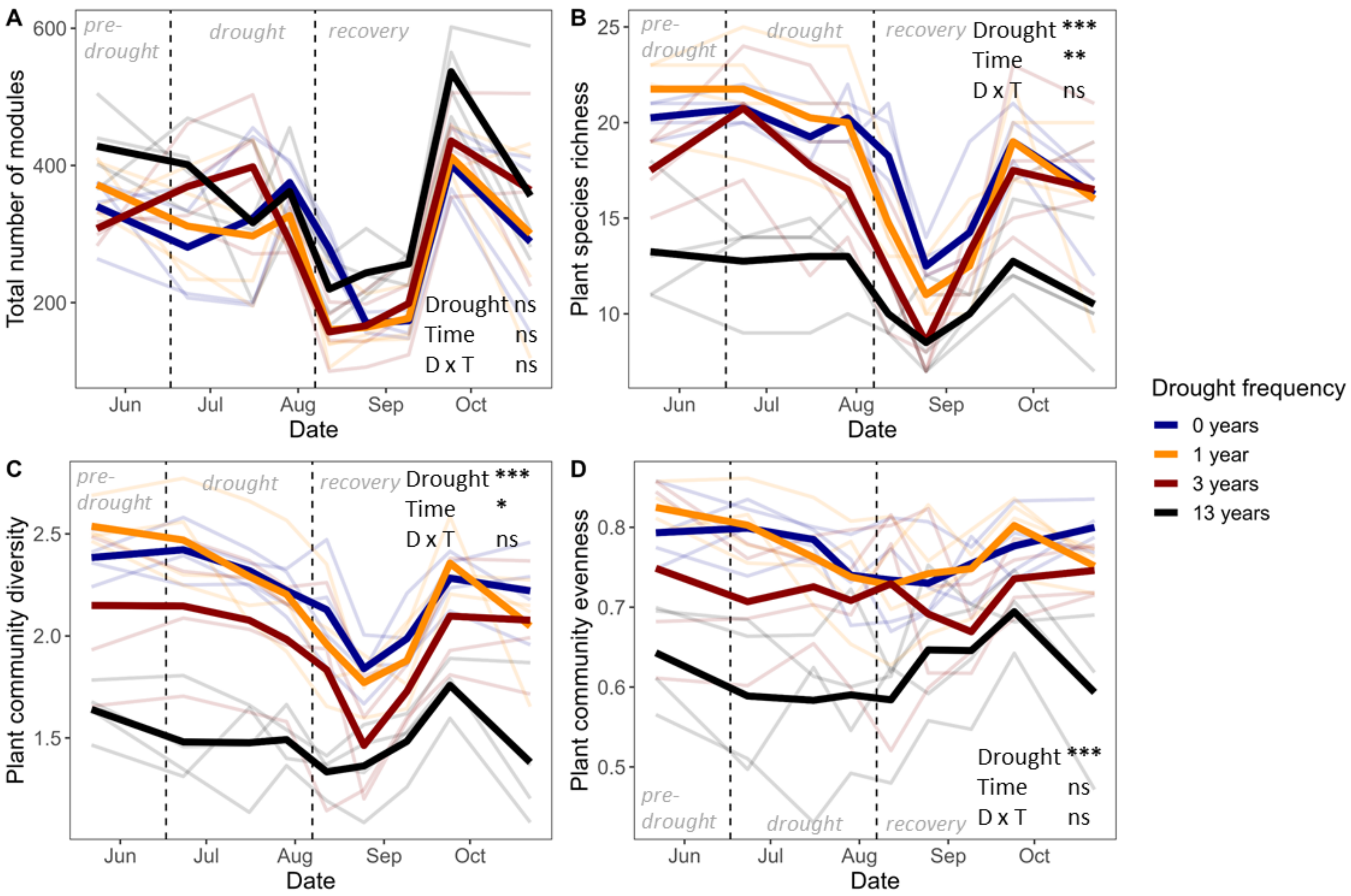


**Fig S6** Drought frequency effects throughout the growing season from mid-May to mid-October on (A) the total number of modules, (B) plant species richness, (C) plant community diversity (Shannon index), and (D) plant community evenness (Pielou's index). Vertical, dotted lines indicate the time points at which drought started (left line) and drought ended, and the plots were mown and rewetted (right line). Solid lines indicate mean numbers, and faded lines indicate numbers of each plot (n = 4 per drought frequency treatment). Results from analyses of variances are shown with '***' indicating p < 0.001, '**' indicating 0.001 > p < 0.01, '*' indicating 0.01 > p < 0.05, and 'ns' indicating no significant effect (p > 0.05).

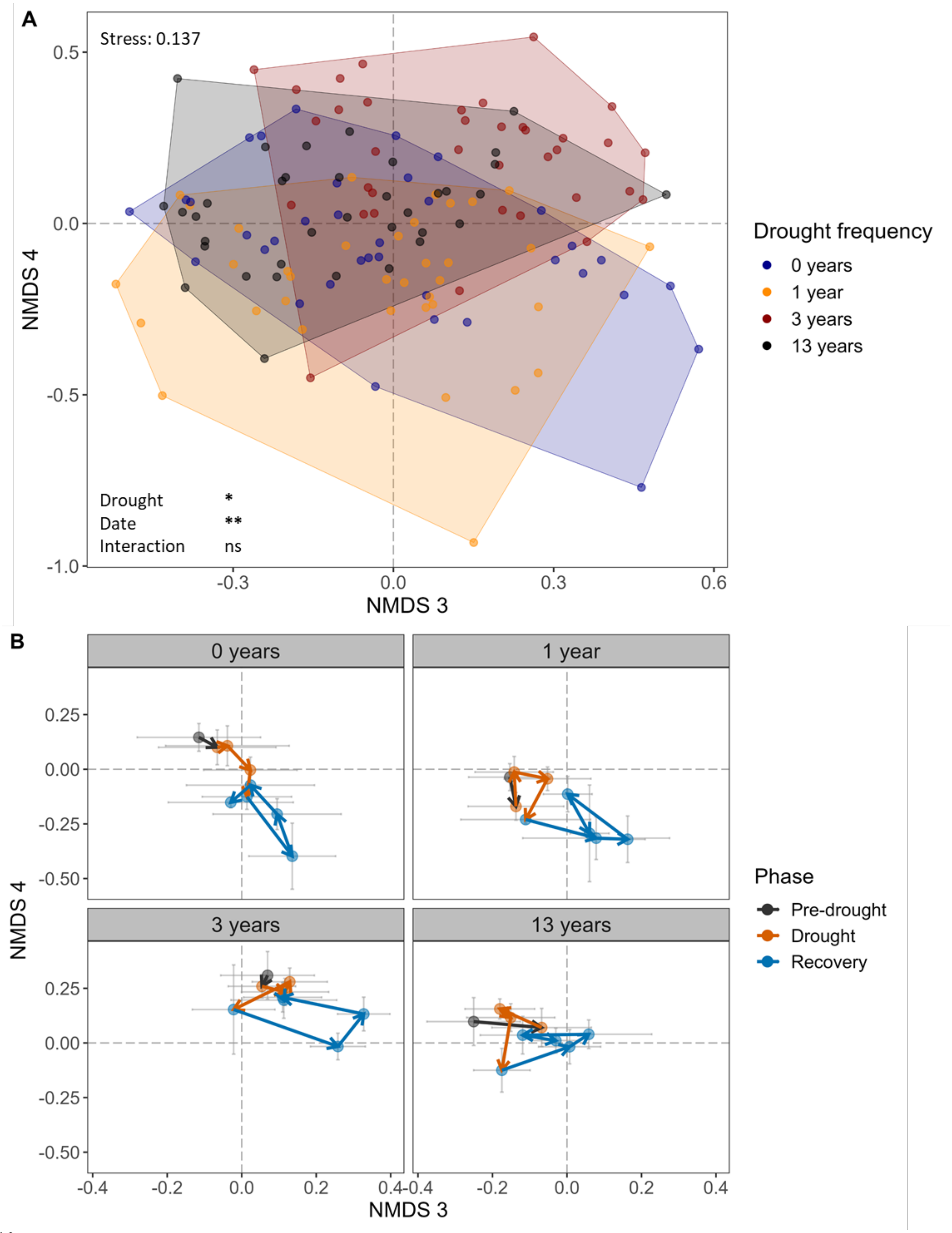


**Fig S7** Non-metric multidimension scaling axes 3 and 4 (NMDS) of plant species module counts throughout the growing season in permanent plots receiving no drought (0 years), one year, three years, and 13 years of recurrent summer drought. (A) Drought frequency effects on plant community composition. Each point represents a plot at a point in time. Hulls group all plots and time points per drought frequency treatment together. Results of a restricted permutation test is shown with ‘**’ indicating $0.001 > p < 0.01$, ‘*’ indicating $0.01 > p < 0.05$, and ‘ns’ indicating no significant effect ($p > 0.05$) ($n = 4$ plots per drought frequency treatment). (B) Community compositional development over time. Points are mean values ± standard errors. Note that axes scales differ between A and B to improve readability. For NMDS axes 1 and 2, see Fig 5. For position of plant species along the axes, see Fig S9.

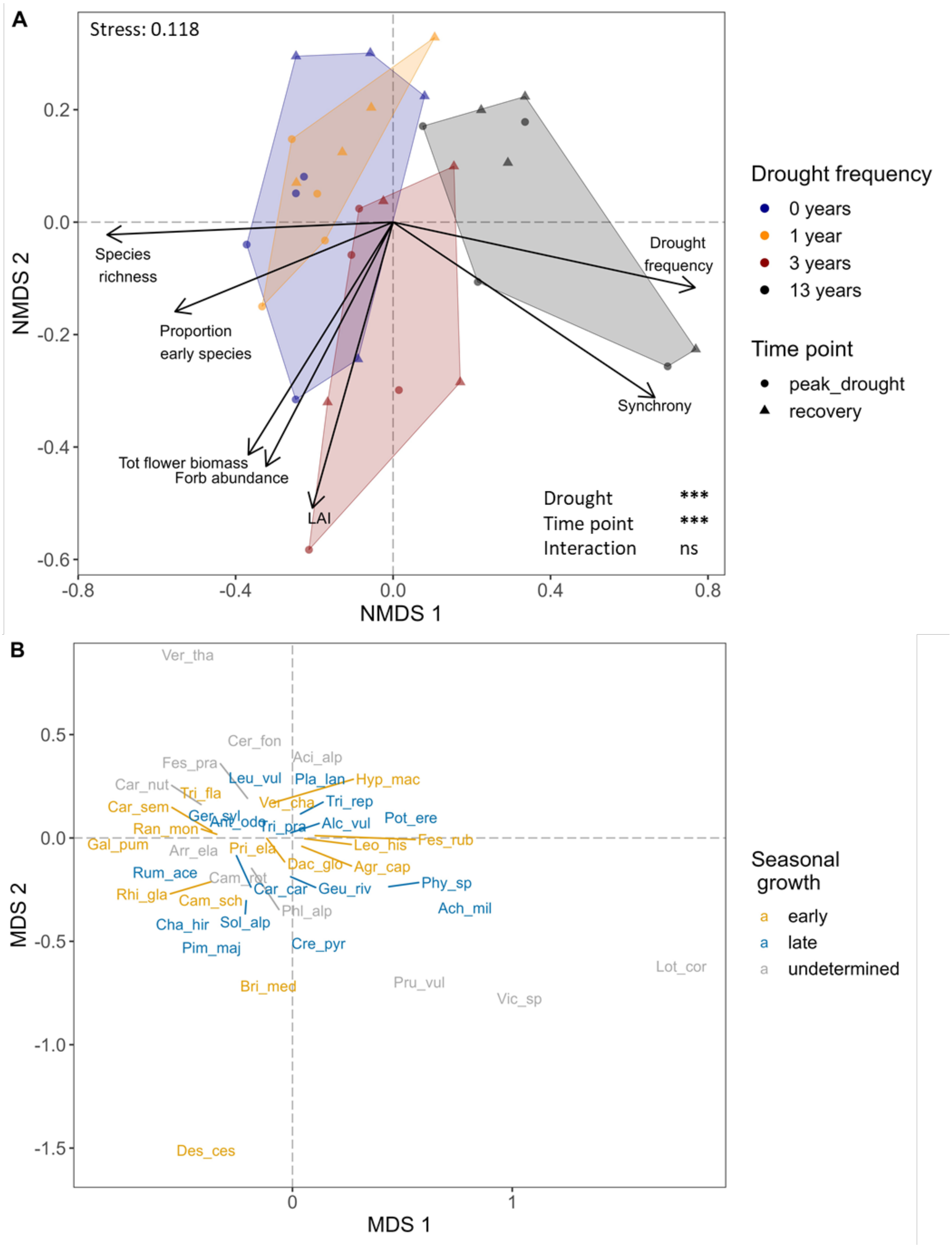


**Fig S8** Non-metric multidimensional scaling (NMDS) of plant community composition based on plant biomass. (A) Plot scores coloured according to drought frequency treatment with shapes indicating the two harvest time points: peak drought (circles) and recovery (triangles). Hulls indicate the area in which all plots per treatment are located. Arrows present an overlay of variables (marginally) significantly relating to the NMDS scores (permutation test with 999 permutations, $p < 0.07$). Results of a restricted permutation test with 999 permutations taking repeated sampling into account are presented. Length of arrows indicate the strength of the relation. ‘Proportion early species’ refers to the proportion of early species versus the total number of species present in a plot. ‘Forb abundance’ refers to the total biomass summed per functional group. ‘Tot flower biomass’ refers to the summed biomass of species flowering stems, and consisted mostly of forb and grass flowers (flower biomass of both functional groups pointed into the same direction as their total biomass, and are not indicated individually to increase readability of the figure). ‘Synchrony’ refers to community seasonal synchrony sensu Loreau & de Mazancourt (2008). ‘LAI’ refers to leaf area index. Results of a restricted permutation test is shown with $p < 0.001$: ***; ns: not significant, $p < 0.05$. (B) Position of the plant species and seasonal growth on the NMDS. Plant species seasonal growth was calculated based on control plots. ‘Early’ seasonal plant species reach their peak number of modules in May-July before the biomass cut. ‘Late’ seasonal plant species reached their peak number of modules in August-October after the biomass cut. ‘Undetermined’ seasonal growth indicates that the species was not present in the control plot and thus seasonal growth could not be determined. For plant species abbreviations, see Table S1.

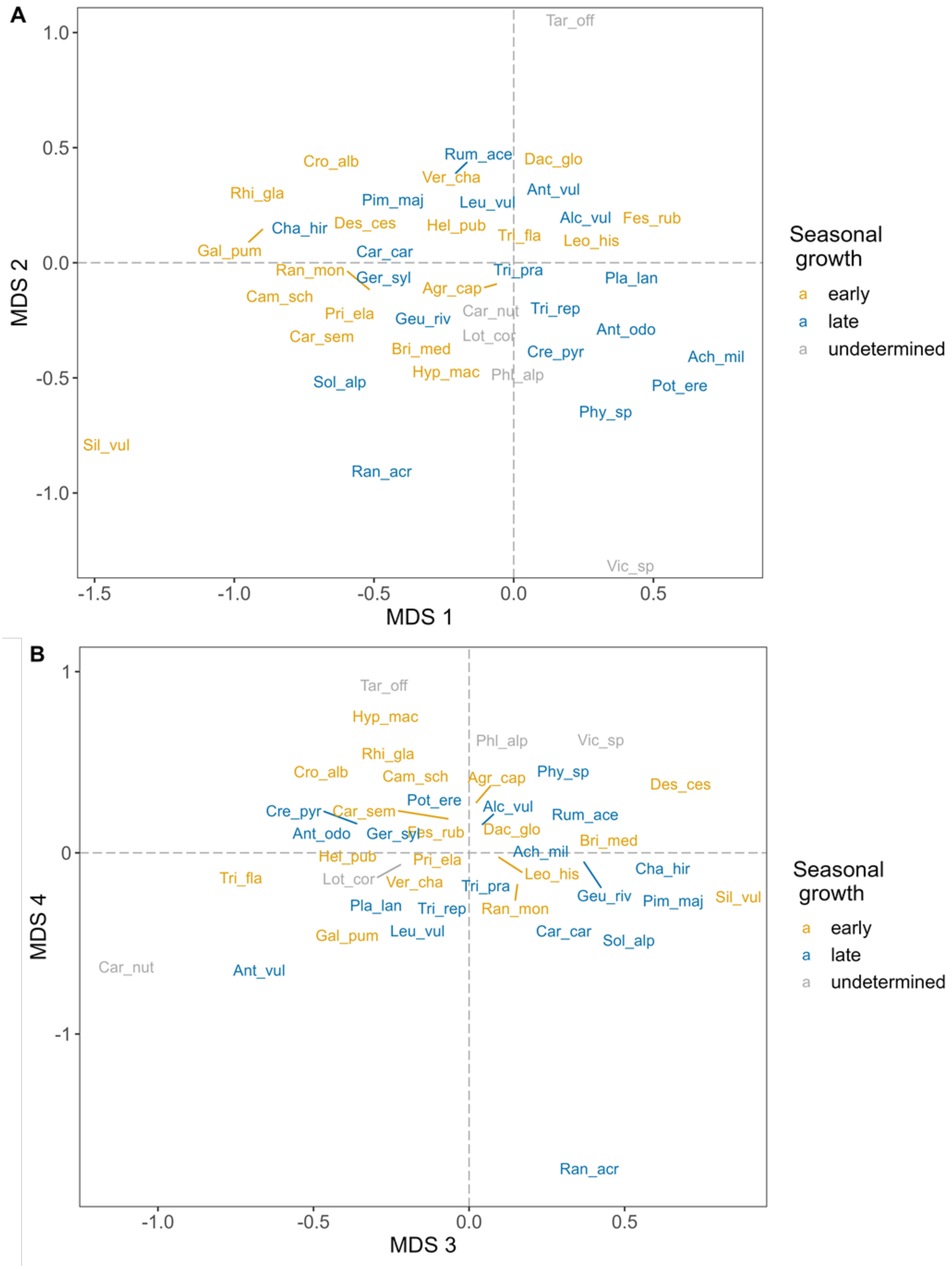


70 **Fig S9** Position of the plant species and their seasonal growth on the NMDS based on module counts. Plant species seasonal growth was calculated based on control plots. 'Early' seasonal plant species

reach their peak number of modules in May-July before the biomass cut. ‘Late’ seasonal plant species reached their peak number of modules in August-October after the biomass cut. ‘Undetermined’ seasonal growth indicates that the species was not present in the control plot and thus seasonal growth could not be determined. For plant species abbreviations, see Table S1. For NMDS site scores, see Fig 5 and Fig S7.

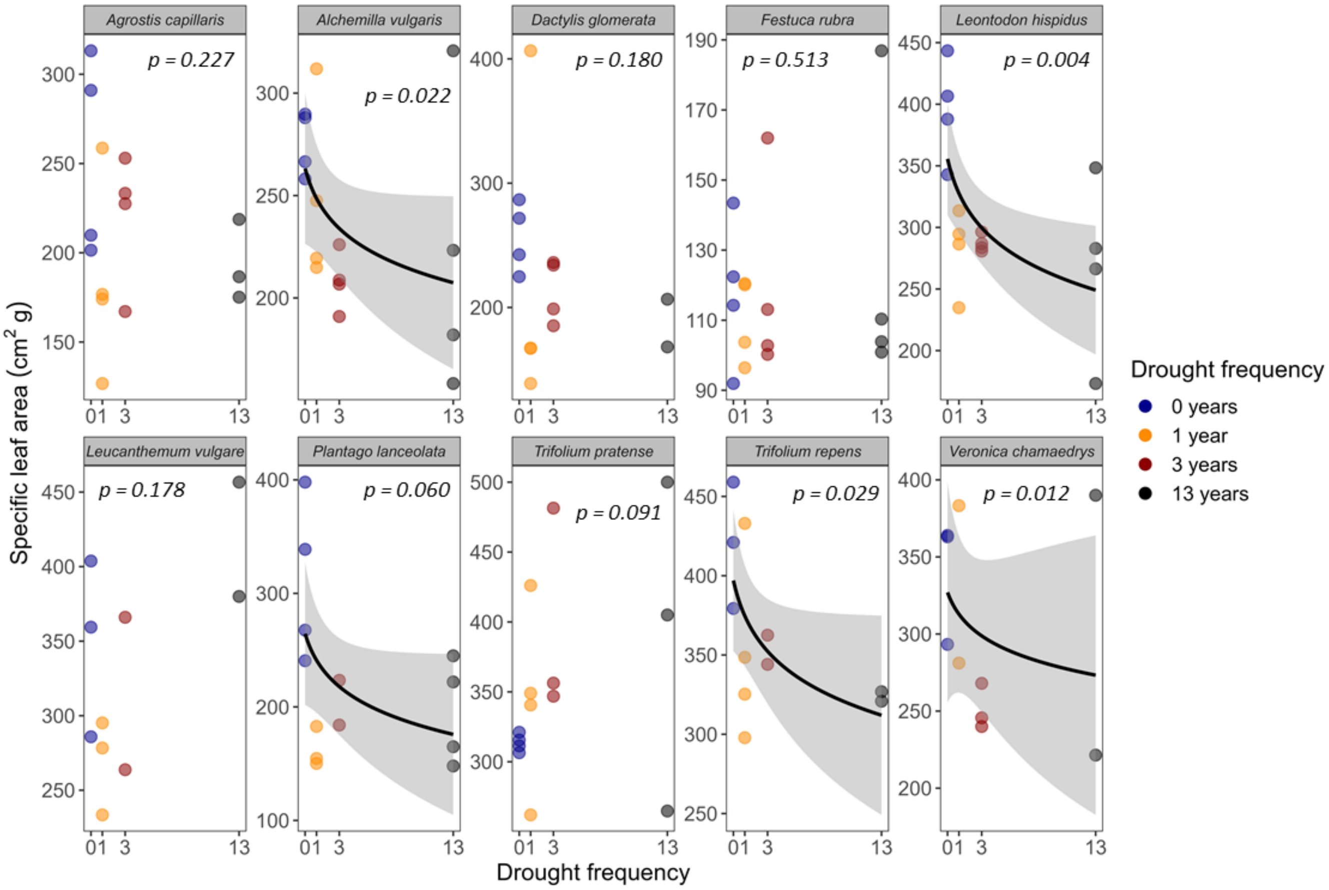


**Fig S10** Drought frequency effects on specific leaf area (SLA) at peak drought of plant species present in all four drought treatments. For significant or marginally significant relations ($p < 0.07$), the mean relation with 95% confidence intervals are shown (n = 10-16 per plant species). For SLA results during the recovery phase, see Fig S11.

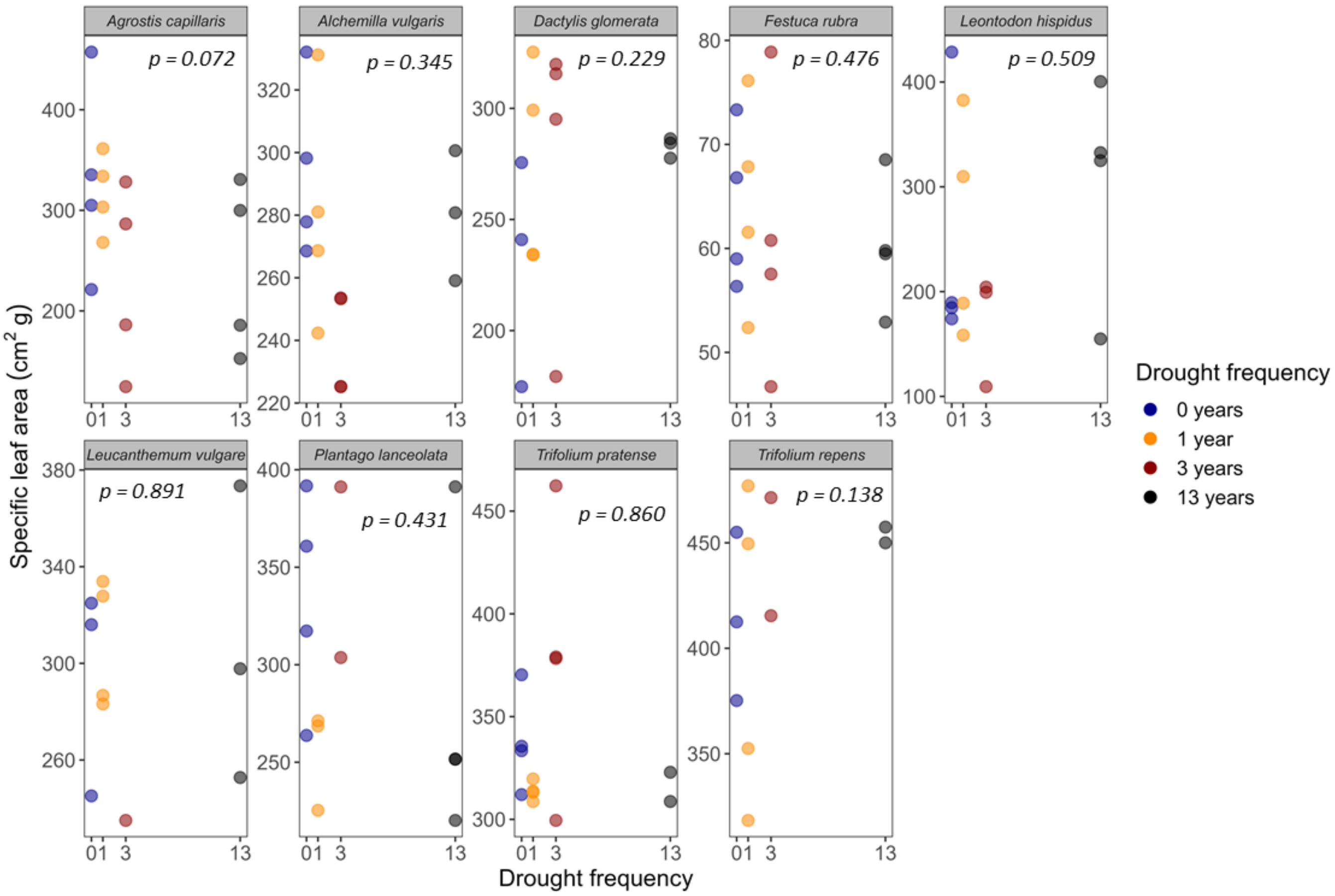


**Fig S11** Drought frequency effects on specific leaf area (SLA) at recovery of plant species present in all four drought treatments. For significant or marginally significant relations ($p < 0.07$), the mean relation with 95% confidence intervals are shown (n = 11-16 per plant species). For SLA results during the peak drought phase, see Fig S10.

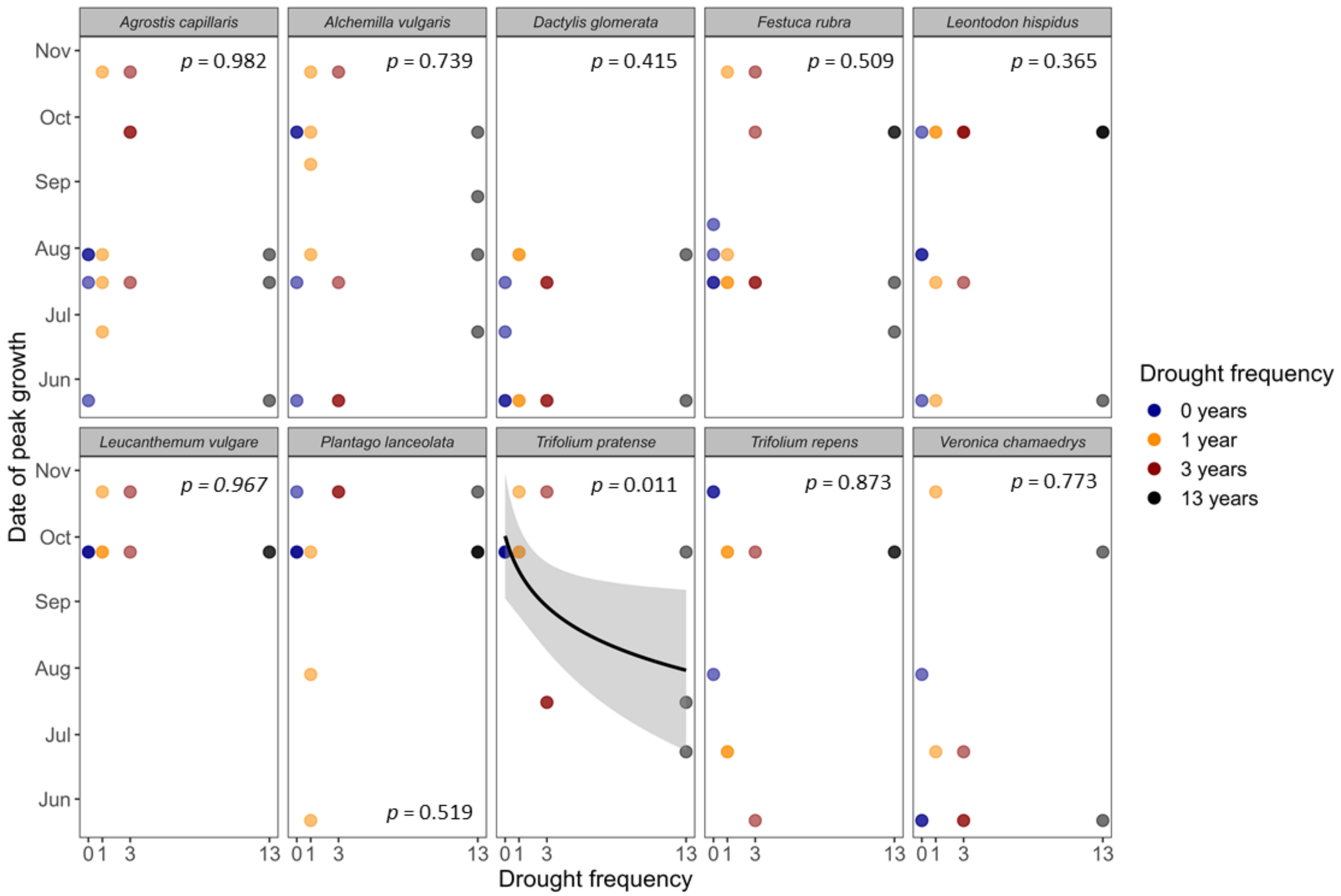


**Fig S12** Drought frequency effects on the timing of seasonal growth of plant species present in all treatments. Seasonal growth was calculated as the date at which the plant species reach its peak number of modules. For significant relations, the mean relation with 95% confidence intervals are shown (n = 10-16 per plant species).

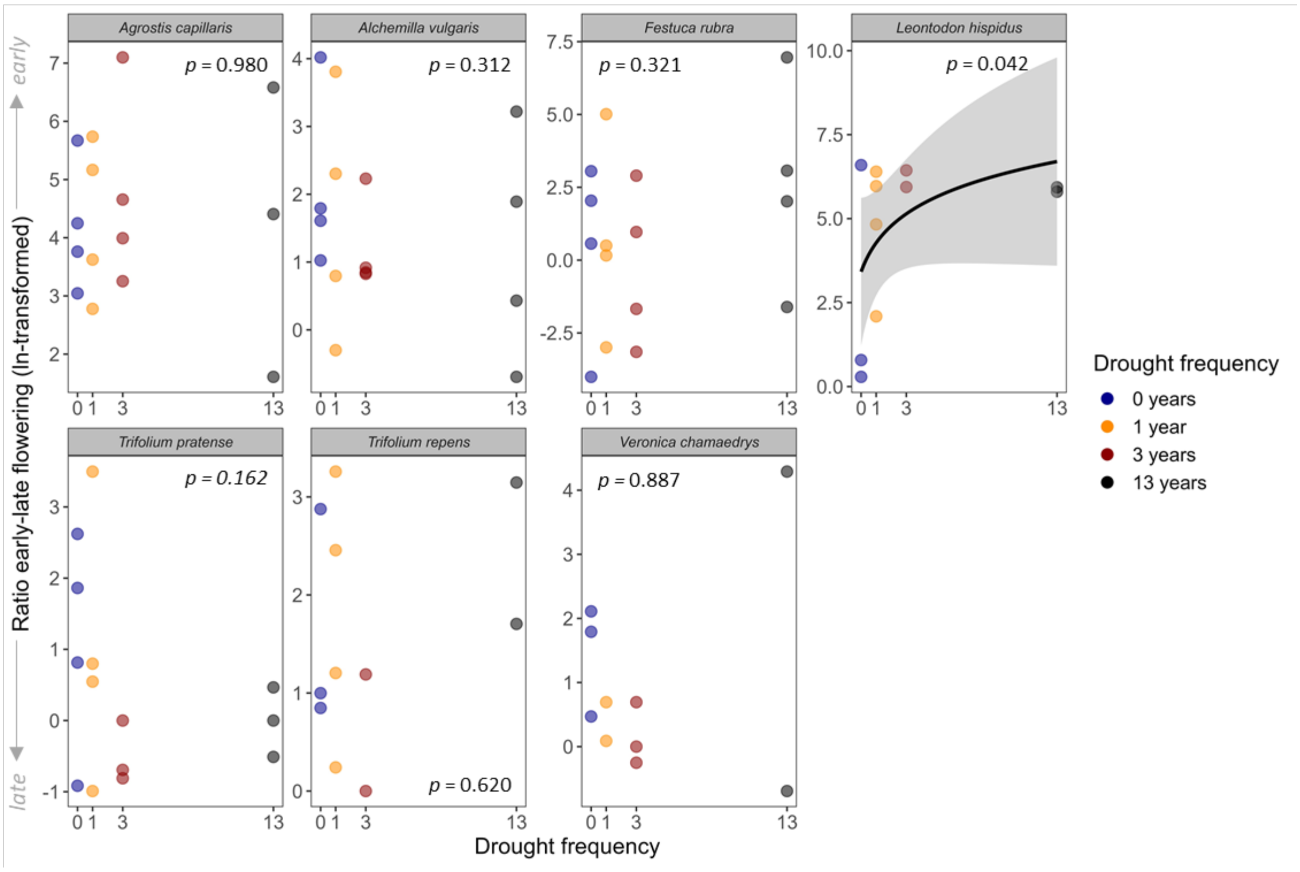


**Fig S13** Drought frequency effects on the timing of flower development of plant species present in all treatments. For significant relations, the mean relation with 95% confidence intervals are shown (n = 10-16 per plant species). A positive value signifies plant species flowering early in the growing season, values near zero indicate species flowering throughout the season, and negative values indicate species that flower later in the growing season.

**Table S1** Names of plant species present in the study site with their respective abbreviations, functional groups, and early/late species specification.

| **Species** | **Abbreviation** | **Functional group** | **Early/Late species** |
|---|---|---|---|
| ***Achillea millefolium*** | Ach_mil | Forb | Late |
| ***Acinos alpinus*** | Aci_alp | Forb | - |
| ***Agrostis capillaris*** | Agr_cap | Grass | Early |
| ***Alchemilla vulgaris*** | Alc_vul | Forb | Late |
| ***Anthoxanthum odoratum*** | Ant_odo | Grass | Late |
| ***Anthyllis vulneraria*** | Ant_vul | Legume | Late |
| ***Arrentherum elatius*** | Arr_ela | Grass | - |
| ***Briza media*** | Bri_med | Grass | Early |
| ***Campanula rotundifolia*** | Cam_rot | Forb | - |
| ***Campanula scheuchzeri*** | Cam_sch | Forb | Early |
| ***Carduus nutans*** | Car_nut | Forb | - |
| ***Carex sempervirens*** | Car_sem | Sedge | Early |
| ***Carum carvi*** | Car_car | Forb | Late |
| ***Cerastium fontanum*** | Cer_fon | Forb | - |
| ***Chaerophyllum hirsutum*** | Cha_hir | Forb | Late |
| ***Crepis pyrenaica*** | Cre_pyr | Forb | Late |
| ***Crocus albiflorus*** | Cro_alb | Forb | Early |
| ***Dactylis glomerata*** | Dac_glo | Grass | Early |
| ***Deschampsia cespitosa*** | Des_ces | Grass | Early |
| ***Festuca pratensis*** | Fes_pra | Grass | - |
| ***Festuca rubra*** | Fes_rub | Grass | Early |
| ***Galium pumilum*** | Gal_pum | Forb | Early |
| ***Geranium sylvaticum*** | Ger_syl | Forb | Late |
| ***Geum rivale*** | Geu_riv | Forb | Late |
| ***Helictotrichon pubescens*** | Hel_pub | Grass | Early |
| ***Hypericum maculatum*** | Hyp_mac | Forb | Early |
| ***Leontodon hispidus*** | Leo_his | Forb | Early |
| ***Leucanthemum vulgare*** | Leu_vul | Forb | Late |
| ***Lotus corniculatus*** | Lot_cor | Legume | - |
| ***Phleum alpinum*** | Phl_alp | Grass | Early |
| ***Phyteuma species (P. orbiculare and P. betonificolia)*** | Phy_sp | Forb | Late |
| ***Pimpinella major*** | Pim_maj | Forb | Late |
| ***Plantago lanceolata*** | Pla_lan | Forb | Late |
| ***Potentilla erecta*** | Pot_ere | Forb | Late |

| | | | |
|---|---|---|---|
| ***Primula elatior*** | Pri_ela | Forb | Early |
| ***Prunella vulgaris*** | Pru_vul | Forb | - |
| ***Ranunculus acris*** | Ran_acr | Forb | Late |
| ***Ranunculus montanus*** | Ran_mon | Forb | Early |
| ***Rhinanthus glacialis*** | Rhi_gla | Hemiparasite | Early |
| ***Rumex acetosa*** | Rum_ace | Forb | Late |
| ***Silene vulgaris*** | Sil_vul | Forb | Early |
| ***Soldanella alpina*** | Sol_alp | Forb | Late |
| ***Taraxacum officinale*** | Tar_off | Forb | - |
| ***Trifolium pratense*** | Tri_pra | Legume | Late |
| ***Trifolium repens*** | Tri_rep | Legume | Late |
| ***Trisetum flavescens*** | Tri_fla | Grass | Early |
| ***Verbascum thapsus*** | Ver_tha | Forb | - |
| ***Veronica chamaedrys*** | Ver_cha | Forb | Early |
| ***Vicia species (V. sepium and V. cracca)*** | Vic_sp | Legume | - |

*‘-‘ in the early-late species column indicates that the species was not present in the control plots to determine its seasonal behaviour.*
